# MRAP2 potentiates GPCR signaling by conserved mechanisms that are disrupted by obesity-associated genetic variants

**DOI:** 10.1101/2025.09.26.678709

**Authors:** Aqfan Jamaluddin, Rachael A. Wyatt, Johannes Broichhagen, Joshua Levitz, Caroline M. Gorvin

**Affiliations:** Department of Metabolism and Systems Science, University of Birmingham, Birmingham, UK; Centre of Membrane Proteins and Receptors (COMPARE), Universities of Birmingham and Nottingham, Birmingham, UK; Department of Biochemistry and Biophysics, Weill Cornell Medicine, New York, NY 10065, USA; Leibniz-Forschungsinstitut für Molekulare Pharmakologie (FMP), 13125, Berlin, Germany

## Abstract

Accessory proteins such as members of the melanocortin-2 receptor accessory protein family (MRAP) have been described to interact with and regulate the signaling of diverse G protein-coupled receptors (GPCRs), however, surprisingly little is known about the mechanisms by which they mediate these effects. MRAP2 modifies signaling of three distinct GPCRs, melanocortin receptor 4 (MC4R), MC3R and the ghrelin receptor (GHSR), which each play essential roles in appetite regulation. Human mutations in MRAP2 cause obesity with hyperglycaemia and hypertension, suggesting that its regulation of GPCRs is critical for maintaining metabolic homeostasis. However, the nature of MRAP2/GPCR complexes and whether there are shared mechanisms for complex assembly, critical structural regions or consistent effects on receptor signaling and trafficking remains unknown. Here we showed all three GPCRs preferentially interact with MRAP2 as 1:1 complexes and that MRAP2 binding disrupts GPCR homodimerization. MRAP2 interacts with the same receptor transmembrane regions to promote GPCR signaling, and the accessory protein impairs β-arrestin-2 recruitment to prolong signaling and delay internalization. Deletion of the cytoplasmic region of MRAP2 impairs GPCR signaling by modulating receptor constitutive activity. Genetic variants in MRAP2 associated with overweight or obesity modulate the constitutive activity of all three GPCRs. Thus, MRAP2 regulates GPCR function using shared molecular mechanisms and these studies provide further evidence of the importance of GHSR constitutive activity.

## Introduction

The hypothalamic leptin-melanocortin axis has an essential role in the central regulation of appetite by integrating information about energy status from the periphery to regulate food intake, energy expenditure and growth. The melanocortin receptor-2 accessory protein-2 (MRAP2) is a single-pass transmembrane protein that is highly expressed in the hypothalamus and modifies the function of several G protein-coupled receptors (GPCRs) that are essential for integrity of the leptin-melanocortin axis^1–4^. MRAP2 was initially described to interact with melanocortin receptor-4 (MC4R)^2^, a GPCR that is stimulated by the biologically active products of pro-opiomelanocortin (POMC) neurons of the hypothalamic arcuate nucleus in the fed-state, and inhibited by AgRP during energy deficit^4^. The central role of MC4R in suppression of food intake has been confirmed in animal studies^5^ and in humans where inactivating MC4R mutations cause hyperphagic, early-onset obesity^6^. MRAP2 also interacts with and facilitates the signaling of another melanocortin receptor family member, MC3R, which is required for the normal activation of AgRP neurons in response to fasting^7,8^, and the growth hormone secretagogue receptor-1a (GHSR), the receptor for ghrelin, a hormone that promotes energy conservation. GHSR activation stimulates AgRP neurons to enhance feeding, promote weight gain and adiposity^9^, activates pituitary somatotrophs to promote growth hormone secretion^10^ and modulates insulin secretion from the pancreas to protect against starvation-induced hypoglycaemia^11^. These actions are suppressed by LEAP2 (liver-expressed antimicrobial peptide-2), an endogenous antagonist and inverse agonist of GHSR which is suppressed by fasting^12,13^. MRAP2 expression on mouse AgRP neurons^1^ and pancreatic δ-cells^14^ are essential for ghrelin-mediated effects on food intake and insulin secretion, respectively. Whether LEAP2 can still suppress GHSR activity in the presence of MRAP2 has not been explored.

It is likely that there are shared mechanisms by which MRAP2 mediates its effects on GPCR signaling, especially for melanocortin receptors. However, most studies focus on a single receptor and there are inconsistencies in the literature driven by the propensity of researchers to overexpress MRAP2 at DNA ratios far exceeding that of its associated GPCR^2,15,16^. Investigations have shown that MRAP2 is still capable of enhancing MC4R and MC3R signaling when expressed at equal concentrations^8,17,18^, and we have shown MC3R forms complexes with MRAP2 with a 1:1 stoichiometry^8^. A preprint has suggested MRAP2 facilitates signaling by disrupting MC4R oligomerization, which normally suppresses receptor signaling^18^, although whether MC4R interacts with MRAP2 monomers or oligomers, which have previously been reported^2,19^, was not explored, nor was disruption of oligomerization examined for other GPCRs. Studies of GHSR have shown MRAP2 effects receptor signaling when expressed at 20x that of GHSR^16^. This suggests GHSR interacts with higher-order oligomers of MRAP2, but no studies have examined how the two proteins interact. A comprehensive examination of complex formation between MRAP2 and GPCRs could reveal whether shared mechanisms exist within receptor subfamilies or across unrelated receptors.

GHSR has considerable basal activity^20^, which is suppressed by GHSR inverse agonists that reduce food intake, body weight, binge-like high fat intake and postprandial blood glucose in animal models^21–23^. GHSR constitutive activity is mediated, at least in part, by the Ala204 residue of extracellular loop-2, which when mutated to Glu204 in humans reduces basal, but not agonist-induced responses, and may contribute to short stature^24^; and in mice, mutation of the equivalent Ala203 residue, impairs ghrelin-mediated food intake, plasma growth hormone, blood glucose and reduces body weight and length in older animals^25^. The Ala204Glu mutation reduces GHSR constitutive activity by hyperpolarizing orexigenic hypothalamic NPY neurons; a mechanism that is shared by LEAP2^25,26^. Cellular studies with high concentrations of MRAP2 suggest MRAP2 likely mediates its physiological effects by reducing GHSR constitutive activity^16^. Whether MRAP2 reduces GHSR constitutive activity by all signaling pathways, and if this is a mechanism shared by other GPCRs remains to be determined.

Modeling of MRAP2 with MC3R and MC4R revealed two structural regions important for MRAP2 effects on receptor signaling. These include interactions between the MRAP2 transmembrane region and TM5-TM6 of the two receptors, and an intracellular helical region that may occlude β-arrestin binding and subsequently enhance signaling by prolonging receptor duration at cell surfaces^8,17^. Impaired agonist-induced recruitment of β-arrestin to MRAP2-coupled GPCRs has been demonstrated for MC4R, MC3R and GHSR^8,16,18^. GHSR has high constitutive internalization^27^, and whether MRAP2 affects either agonist-dependent or -independent GHSR trafficking or recruitment of β-arrestin under basal conditions is unknown. Investigation of the role of the MRAP2 cytoplasmic region in β-arrestin recruitment and other functions (e.g. G protein recruitment) for multiple GPCRs may provide further insights into the role of this structural region. Moreover, whether TM5 and TM6 are essential for other GPCR interactions with MRAP2 or just melanocortin receptors warrants investigation.

Mice with deletion of the MRAP2 gene develop extreme obesity, visceral adiposity and increased fat mass, similar to MC4R knockout mice^5,28^. However, MRAP2-depleted mice lack the early-onset hyperphagia of MC4R knockout mice, which has been attributed to the absence of MRAP2 effects on GHSR^1^. Genetic variants in MRAP2, which impair signaling by MC4R, are associated with overweight and obesity in humans^8,17,29^. Hyperglycaemia, hypertension and high blood cholesterol are more frequently observed in these individuals than those with MC4R mutations^29^, which could suggest these MRAP2 variants affect activity of other GPCRs including GHSR. Genetic variants that affect GHSR signaling have not consistently been linked with human disease. Associations were identified between predicted loss-of-function variants in GHSR and standing height in the UK Biobank^30^, and several studies indicate that impaired constitutive activity correlates with short stature^24,31^, with some evidence of lower BMI in early life^32,33^. Whether obesity-associated MRAP2 variants affect GHSR constitutive activity or agonist-induced signaling remains to be explored.

Here we performed a comprehensive analysis of the mechanisms by which MRAP2 enhances signaling by GHSR, MC3R and MC4R. Our studies provide evidence that MRAP2 disrupts GPCR oligomerization to interact with monomeric GPCRs and that the MRAP2 cytoplasmic region has an essential role in agonist-mediated G protein signaling by multiple GPCRs. Human MRAP2 variants enhance GHSR constitutive activity and internalization, which may contribute to increased weight gain and hyperglycaemia observed in individuals with these genetic variants.

## Results

### GHSR interacts with MRAP2 monomers primarily at a 1:1 ratio

Previous studies using coimmunoprecipitation and NanoBiT have shown GHSR interacts with MRAP2, although MRAP2 was expressed at 5-20 times that of GHSR to observe a signaling response^1,16^. Our previous studies have shown MRAP2 interacts with MC3R in a 1-to-1 stoichiometry and that high concentrations of MRAP2 are not necessary to observe changes in signaling^8^. To assess the stoichiometry with which GHSR interacts with MRAP2 we used the single-molecule pull-down (SiMPull) technique^34^ (Figure 1A). MRAP2 constructs with N-terminal HA or FLAG epitopes and SNAP, Halo or CLIP tags have been described and shown to enhance signaling by MC4R and MC3R^8,17^. We generated GHSR constructs with N-terminal hemagglutinin (HA) or FLAG epitopes followed by a SNAP, Halo or CLIP tag amenable to labelling with organic dyes (Figure S1A)^8,17^. GHSR plasmids signaled similarly to untagged GHSR and commercially available plasmids (SNAP-GHSR) (Figure S1B-C), trafficked normally (Figure S1D-E) and colocalized with MRAP2 when transiently transfected in HEK293 cells (Figure S1F). We first used SiMPull to examine the stoichiometry of GHSR homomers. Cells were transfected with HA-Halo-GHSR and labelled with membrane impermeable CA-Sulfo646^35^, then cells were lysed, receptors immobilized by anti-HA antibodies and single molecules imaged by total internal reflection fluorescence microscopy (Figure 1B). The majority of GHSR molecules (66%) showed single bleaching steps (i.e. monomers), although GHSR also forms dimers (31%) and occasionally higher-order oligomers with three or four photobleaching steps at the cell surface (Figure 1C-D). In the absence of anti-HA tagged protein, few molecules were observed (105 molecules across 10 images) (Figure 1E). Therefore, GHSR primarily forms stable monomers or dimers when expressed alone.

**Figure 1.**
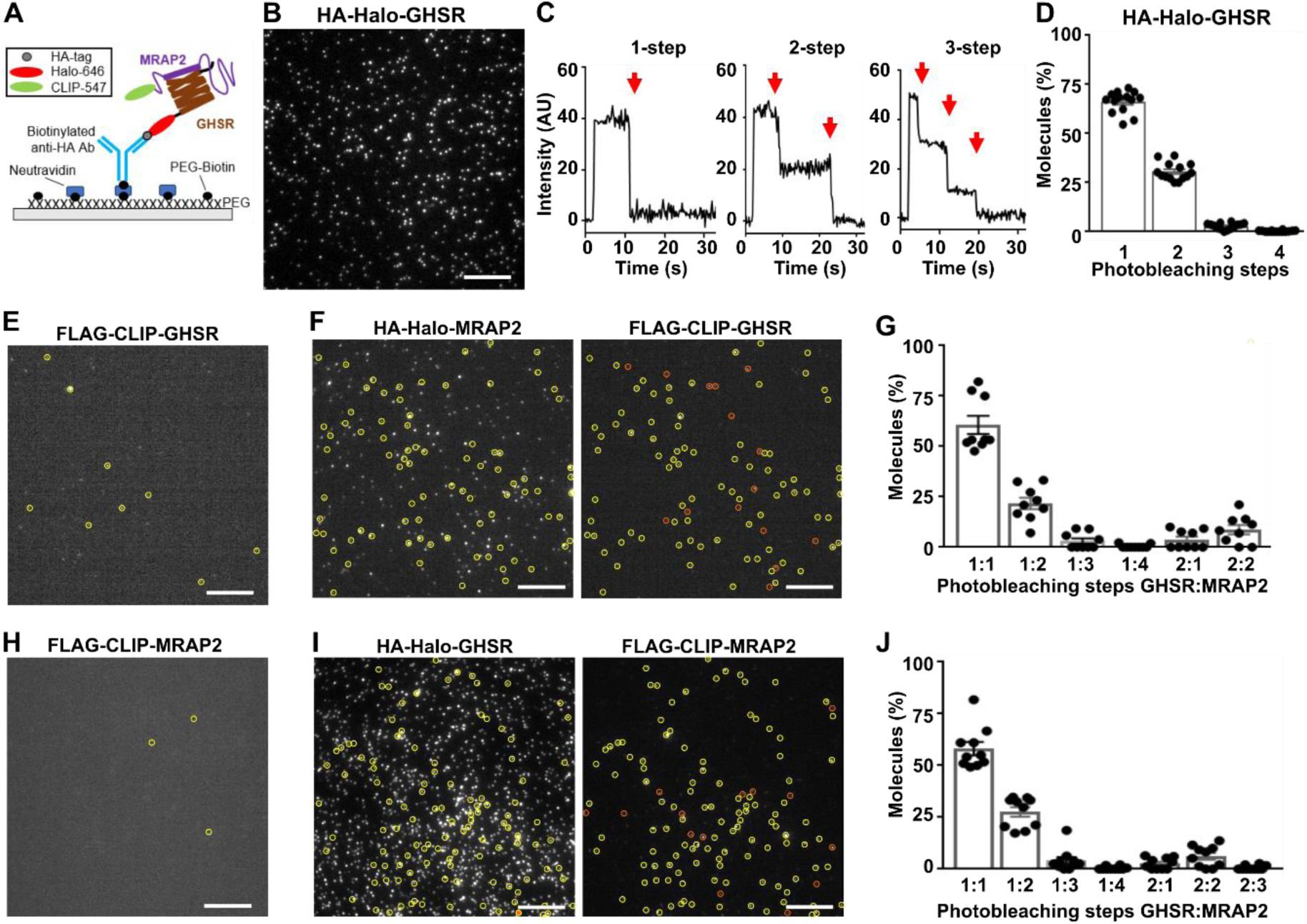
GHSR interacts with MRAP2 monomers primarily at a 1:1 ratio. (**A**) Cartoon showing two-color SiMPull experiments. Fresh cell lysate from HEK293 cells co-expressing HA-Halo-GHSR and FLAG-CLIP-MRAP2 is added to a PEG-passivated glass slide with immobilized anti-HA antibody. Halo and CLIP tags are labeled with CA-Sulfo646 and BC-DY547, respectively. (**B**) Representative single-molecule fluorescence image of HA-Halo-GHSR with (**C**) examples of single-molecule fluorescence traces with photobleaching steps indicated with red arrows. (**D**) Proportion of molecules with 1 to 4 bleaching steps. N = 3240 molecules from 12 movies. (**E**) Cells transfected with FLAG-CLIP-GHSR only, showing negligible background fluorescence. (**F**) Representative two-color SiMPull images of HA-Halo-MRAP2 and FLAG-CLIP-GHSR with colocalized spots circled in yellow and non-colocalized spots shown in orange. (**G**) Photobleaching step analysis from colocalized spots. N = 872 from 9 movies. (**H**) Cells transfected with FLAG-CLIP-MRAP2 only, showing negligible background fluorescence. (**I**) Representative two-color SiMPull images of HA-Halo-GHSR and FLAG-CLIP-MRAP2 with colocalized spots circled in yellow, and (**J**) Photobleaching step analysis from colocalized spots. N = 953 from 10 movies. Scale, 10 μm. See also Figure S2.

To investigate GHSR and MRAP2 heteromers, HA-Halo-MRAP2 and FLAG-CLIP-GHSR were transfected in HEK293 cells and Halo and CLIP tags labelled with CA-Sulfo646 and BC-DY547 fluorophores, respectively, then cells were lysed, receptors immobilized by anti-HA antibodies and fluorescence co-localization assessed. Co-localization was present in 22% of GHSR spots (Figure S2A). Photobleaching step analysis showed 1-step each for GHSR and MRAP2 in ∼61% of co-localized spots, while 2-step bleaching for MRAP2 was observed in 22% (Figure 1F-G). Almost 9% of spots showed two GHSR and two MRAP2 bleaching steps. SiMPull experiments were repeated with the Halo and CLIP labels swapped. Cells were co-transfected with HA-Halo-GHSR and FLAG-CLIP-MRAP2, then labelled and imaged. FLAG-CLIP-MRAP2 expression alone produced few single molecules (Figure 1H), as previously described^8^, demonstrating the pull-down is specific to HA-tagged proteins. Co-localization was present in ∼27% of MRAP2 spots (Figure S2B). Photobleaching step analysis showed 58% had one GHSR and one MRAP2 step, while 2-step bleaching for MRAP2 was observed in 27% of spots, and 6% had two steps each for GHSR and MRAP2 (Figure 1I-J). Therefore, GHSR is more likely to interact with MRAP2 in a 1:1 stoichiometry but can interact with more than one MRAP2 molecule.

### Dimerization with MRAP2 disrupts GPCR oligomerization

A recent preprint has indicated that MRAP2 may enhance MC4R signaling by disrupting GPCR oligomerization^18^. We sought to explore these findings in more detail and assess whether GHSR and other receptors may share this mechanism using SiMPull. We first examined the stoichiometry of MC4R homomers in cells transfected with HA-Halo-MC4R and labelled with membrane impermeable CA-Sulfo646. Cells were lysed, receptors immobilized by anti-HA antibodies and single molecules imaged (Figure 2A). Approximately 64% of cell surface MC4R molecules showed single bleaching steps, while 31% were dimers and almost 5% had three photobleaching steps (Figure 2B). Therefore, MC4R is primarily expressed as a monomer at the cell surface, although one-third of molecules form dimers or higher-order oligomers, similarly to GHSR.

**Figure 2.**
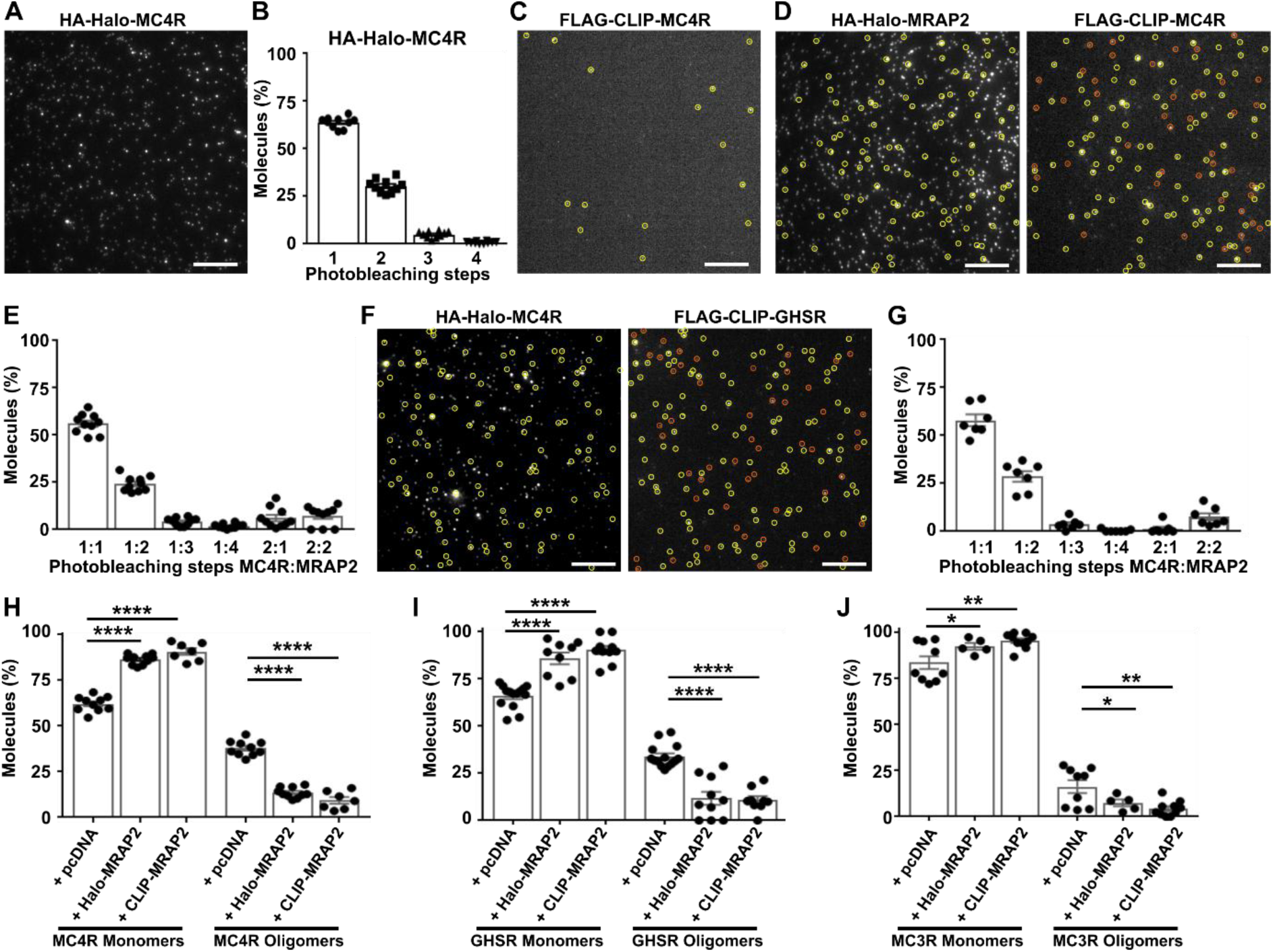
Dimerization with MRAP2 disrupts GPCR oligomerization. (**A**) Representative single-molecule fluorescence image of HA-Halo-MC4R with (**B**) quantification of the proportion of molecules with 1 to 4 bleaching steps. N = 3550 molecules from 10 movies. (**C**) Cells transfected with FLAG-CLIP-MC4R only, showing negligible background fluorescence. (**D**) Representative two-color SiMPull images of HA-Halo-MRAP2 and FLAG-CLIP-MC4R with colocalized spots circled in yellow and non-colocalized spots shown in orange. (**E**) Photobleaching step analysis from colocalized spots. N = 1249 from 10 movies. (**F**) Representative two-color SiMPull images of HA-Halo-MC4R and FLAG-CLIP-MRAP2 with colocalized spots circled in yellow and non-colocalized spots shown in orange. (**G**) Photobleaching step analysis from colocalized spots. N = 989 from 7 movies. Scale, 10 μm for all. (**H-J**) Quantification of monomeric vs. oligomeric GPCR in single-molecule spots from cells co-expressing pcDNA, Halo-MRAP2 or CLIP-MRAP2 and (H) MC4R, (I) GHSR, and (J) MC3R. Statistical analyses were performed by one-way ANOVA with Tukey’s multiple comparisons test. ****p<0.0001, ***p<0.001, **p<0.01, *p<0.05. See also Figure S2.

HEK293 cells were then co-transfected with HA-Halo-MRAP2 and FLAG-CLIP-MC4R, labelled and SiMPull analyses performed to investigate MC4R and MRAP2 complexes. FLAG-CLIP-MC4R expression alone produced few single molecules (Figure 2C). Co-localization was present in 29% of MC4R spots (Figure S2C). Photobleaching step analysis showed 1-step each for MC4R and MRAP2 in ∼56% of co-localized spots, while 2-step bleaching for MRAP2 was observed in 24%, 2-step bleaching for MC4R was observed in 6%, and 7% of spots showed two MC4R and two MRAP2 bleaching steps (Figure 2D-E). Similar results were obtained when the Halo and CLIP labels were swapped. Co-localization was present in ∼28% of MRAP2 spots when cells were co-transfected with HA-Halo-MC4R and FLAG-CLIP-MRAP2 (Figure S2D). Photobleaching step analysis showed 58% had one MC4R and one MRAP2 step, while 2-step bleaching for MRAP2 was observed in 29% of spots, and almost 8% had two steps each for MC4R and MRAP2 (Figure 2F-G). Therefore, similarly to GHSR, MC4R is more likely to interact with MRAP2 in a 1:1 stoichiometry but can interact with more than one MRAP2 molecule.

To determine whether MRAP2 disrupts the proportion of oligomeric GPCRs, two-color SiMPull data was reanalyzed to assess the number of GPCRs in any monomeric form (e.g. 1:1, 1:2, 1:3, 1:4 GPCR to MRAP2) compared to the number of receptors in oligomeric form (e.g. 2:1, 2:2 GPCR to MRAP2). Analyses were first performed on data from cells co-expressing MC4R and MRAP2. This showed a significantly higher proportion of monomeric MC4R when co-expressed with Halo-MRAP2 (∼86%) or CLIP-MRAP2 (∼91%) than in cells expressing MC4R and pcDNA (∼62%) (Figure 2H). Data from cells co-expressing GHSR and MRAP2 also showed a significantly higher proportion of monomeric GHSR when co-expressed with Halo-MRAP2 (∼86%) or CLIP-MRAP2 (∼90%) than in cells expressing GHSR and pcDNA (∼66%) (Figure 2I). Finally, we analyzed our previous datasets in which we examined MC3R colocalization with MRAP2 by SiMPull^8^. MC3R had less propensity to oligomerize than did MC4R or GHSR. However, a significantly higher proportion of monomeric MC3R was still observed when the receptor was co-expressed with Halo-MRAP2 (∼96%) or CLIP-MRAP2 (∼93%) than in cells expressing MC3R and pcDNA (∼84%) (Figure 2J). This was not due to a reduction in the total number of GPCR molecules at the cell surface as MRAP2 co-expression increased the amount of GHSR at the cell surface (Figure S2E-F) or had no effect on total surface expression of MC3R and MC4R (shown in previous analyses^8,17^). Therefore, disruption of GPCR oligomers by MRAP2 may be a general mechanism by which it interacts with receptors.

### MRAP2 enhances GHSR-Gq and Gi signaling and reduces internalization

Previous studies have shown that MRAP2 enhances GHSR-IP3 signaling by suppressing receptor constitutive activity and reducing β-arrestin recruitment when MRAP2 is expressed at 5-20x GHSR concentrations. We first sought to assess whether MRAP2 could facilitate GHSR at 1:1 ratios with low receptor expression (50 ng of each receptor). HEK293 cells were transfected with HA-SNAP-GHSR and pcDNA or MRAP2-WT then IP3 activation assessed by NanoBiT biosensor^36^ and intracellular calcium (Ca^2+^_i_) accumulation measured by Fluo-4 assay. In response to increasing concentrations of the GHSR agonist MK-0677, MRAP2 significantly enhanced IP3 and Ca^2+^_i_ maximal responses and reduced constitutive activity but had no effect on pEC50 values (Figure 3A-B, Table S2). MRAP2 enhances IP3-Ca^2+^_i_ by increasing recruitment of mGq (assessed by BRET) under basal and agonist-driven conditions in both HEK293 and the mHypo-N39 mouse hypothalamic cell-line (Figure 3C, Figure S3A, Table S2). GHSR can also couple to Gi/o signaling pathways^37^ and to assess this pathway cells were pre-incubated with forskolin to increase cAMP concentrations, then GHSR-mediated suppression of cAMP measured in the presence of pcDNA or MRAP2. Co-expression of MRAP2 with GHSR significantly enhanced Gi/o-mediated suppression of forskolin-induced cAMP (Figure 3D, Table S2).

**Figure 3.**
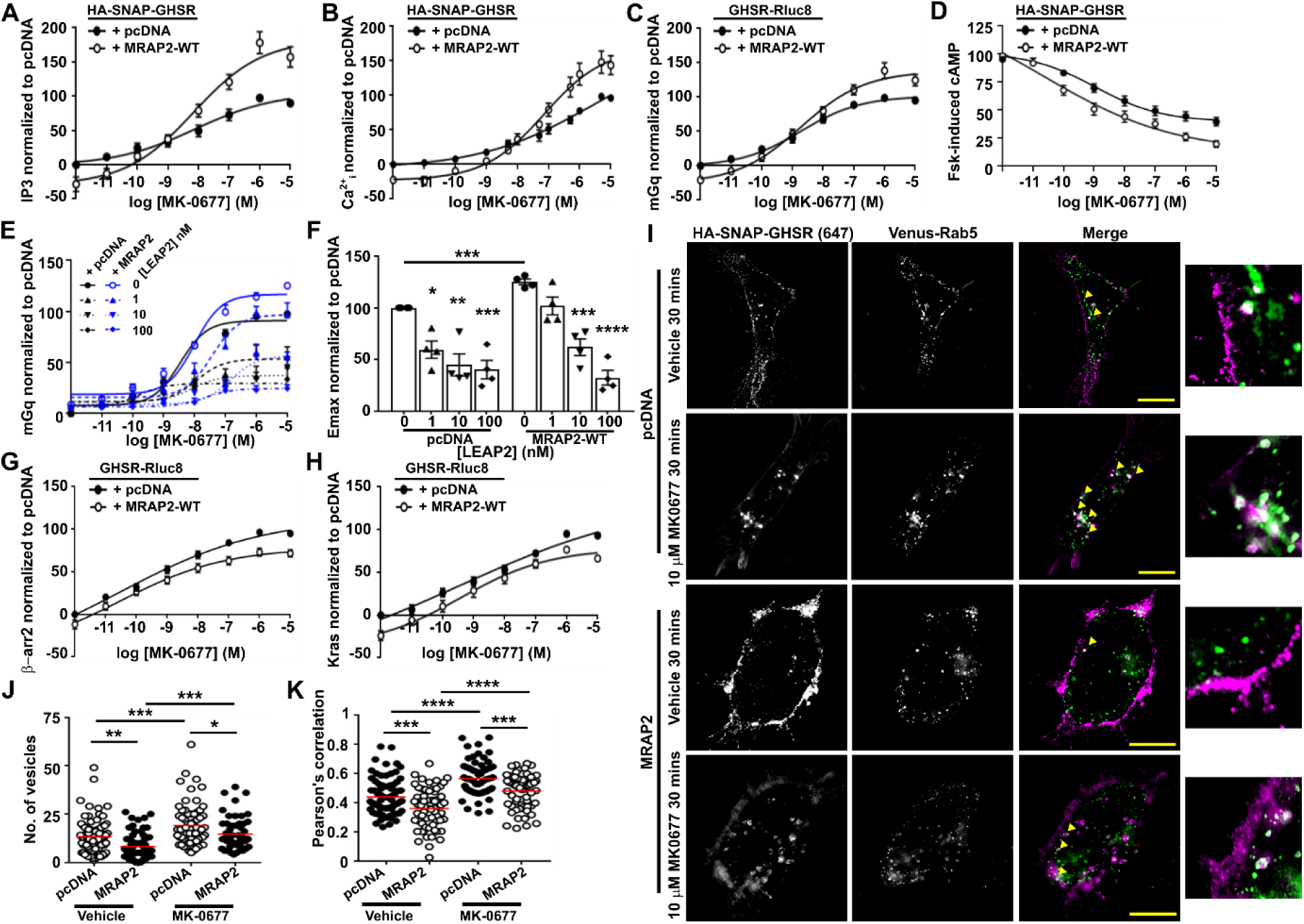
MRAP2 enhances GHSR Gq and Gi signaling and reduces internalization. Cells were transfected with either pcDNA, MRAP2 WT or MRAP2 mutant proteins and the following GHSR-mediated responses assessed: (**A**) Normalized IP3 measured by IP3 NanoBiT (N = 11), (**B**) Normalized intracellular calcium (Ca^2+^_i_) measured by Fluo-4 (N = 9), (**C**) Normalized BRET between GHSR-Rluc8 and mGq (N = 10), (**D**) suppression of forskolin-induced cAMP responses (N = 12), (**E**) Normalized BRET between GHSR-Rluc8 and mGq in cells pre-incubated with the GHSR antagonist LEAP2, with (**F**) Emax responses (N = 4). (**G-H**) Normalized BRET between GHSR-Rluc8 and (G) β-arrestin-2-YFP and (H) Venus-Kras (N = 10 in G, N = 13 in H). AUC was measured in A, C, D, E, G, H and responses expressed relative to the pcDNA maximal response set at 100. Data shows mean±SEM in A-H. Effects on mGq and β-arrestin-2 recruitment and Kras dissociation were replicable in mHypo-N39 mouse hypothalamic cell-lines (Figure S3). (**I**) SIM imaging of GHSR and Venus-Rab5 in the presence of pcDNA or MRAP2 with (right) closer view, with (**J**) quantification of the number of internalized vesicles and (**K**) correlation between GHSR and Rab5 assessed by Pearson’s coefficient. Yellow arrows show colocalization. Scale, 5 μm. N = 63-73 cells from seven independent transfections for each group in J and K. Red line shows mean in J and K. Statistical analyses were performed by one-way ANOVA with Dunnett’s multiple-comparisons test in E and by one-way ANOVA with Sidak’s multiple-comparisons test in J and K. Statistics for constitutive activity, Emax and pEC50 for other panels are shown in Table S2 or S3.****p<0.0001, ***p<0.001, **p<0.01, *p<0.05.

LEAP2 is an endogenous antagonist of GHSR, although whether it still binds to the receptor in the presence of MRAP2 has not been determined^12,13^. mGq recruitment was assessed by BRET in the presence of four concentrations of LEAP2 in cells transfected with GHSR-Rluc8 and pcDNA or MRAP2. Exposure of cells to 1, 10 and 100 nM LEAP2 reduced GHSR-mediated mGq activity and Emax, such that MRAP2-mediated signaling was not significantly different to cells expressing pcDNA and GHSR (Figure 3E-F). LEAP2 did not affect pEC50 values (Table S3), consistent with previous studies^38^.

MRAP2 has been described to reduce the ability of several GPCRs, including GHSR, to recruit β-arrestin^8,16^. We confirmed that MRAP2 reduces the recruitment of β-arrestin-2 to GHSR when compared to cells co-transfected with pcDNA, and this was associated with an increase in pEC50 (Figure 3G, Table S2). This was replicated in mHypo-N39 mouse hypothalamic cells (Figure S3B). MRAP2 also reduces GHSR internalization, assessed by BRET dissociation from the plasma membrane marker Kras in AdHEK and mHypo-N39 (Figure 3H, Figure S3C, Table S2), and significantly delays recruitment to Rab5-positive endosomes (Figure 3I-K). MRAP2 reduced both constitutive and agonist-driven internalization. This suggests MRAP2 enhances signaling by suppressing receptor constitutive activity and impairing β-arrestin recruitment.

### GHSR TM5-TM7 are important for GHSR-MRAP2 interactions and MRAP2-mediated signaling

We next sought to determine the transmembrane structural regions that are involved in GHSR and MRAP2 interactions. We designed transmembrane (TM) interference peptides to mimic the sequence of each GHSR TM helix. These TM interfering peptides were fused to the cell-penetrating TAT sequence, and we first tested their ability to disrupt GHSR-MRAP2 complex formation in a NanoBiT luminescence complementation assay. LgBiT was cloned to the C-terminus of GHSR (GHSR-LgC) and transfected with either the previously described MRAP2-SmC^8^ or the SmC empty vector (negative). Luminescence was measured in comparison to cells transfected with LgC and SmC empty vectors as a negative control. Luminescence values were 400-fold higher in cells transfected with GHSR and MRAP2 than those with negative controls (Figure 4A). NanoBiT assays were then repeated with the TM interfering peptides. Addition of peptides targeting all GHSR TM helices simultaneously abolished GHSR-MRAP2 complex formation such that luminescence values were indistinguishable to those of the negative control. Peptides replicating TM5, TM6 and TM7 significantly reduced NanoBiT luminescence, indicating the involvement of their interfaces in formation of the GHSR-MRAP2 complex (Figure 4B). In contrast, TM1-TM4 mimicking peptides had no effect on GHSR-MRAP2 complex formation (Figure 4B).

**Figure 4.**
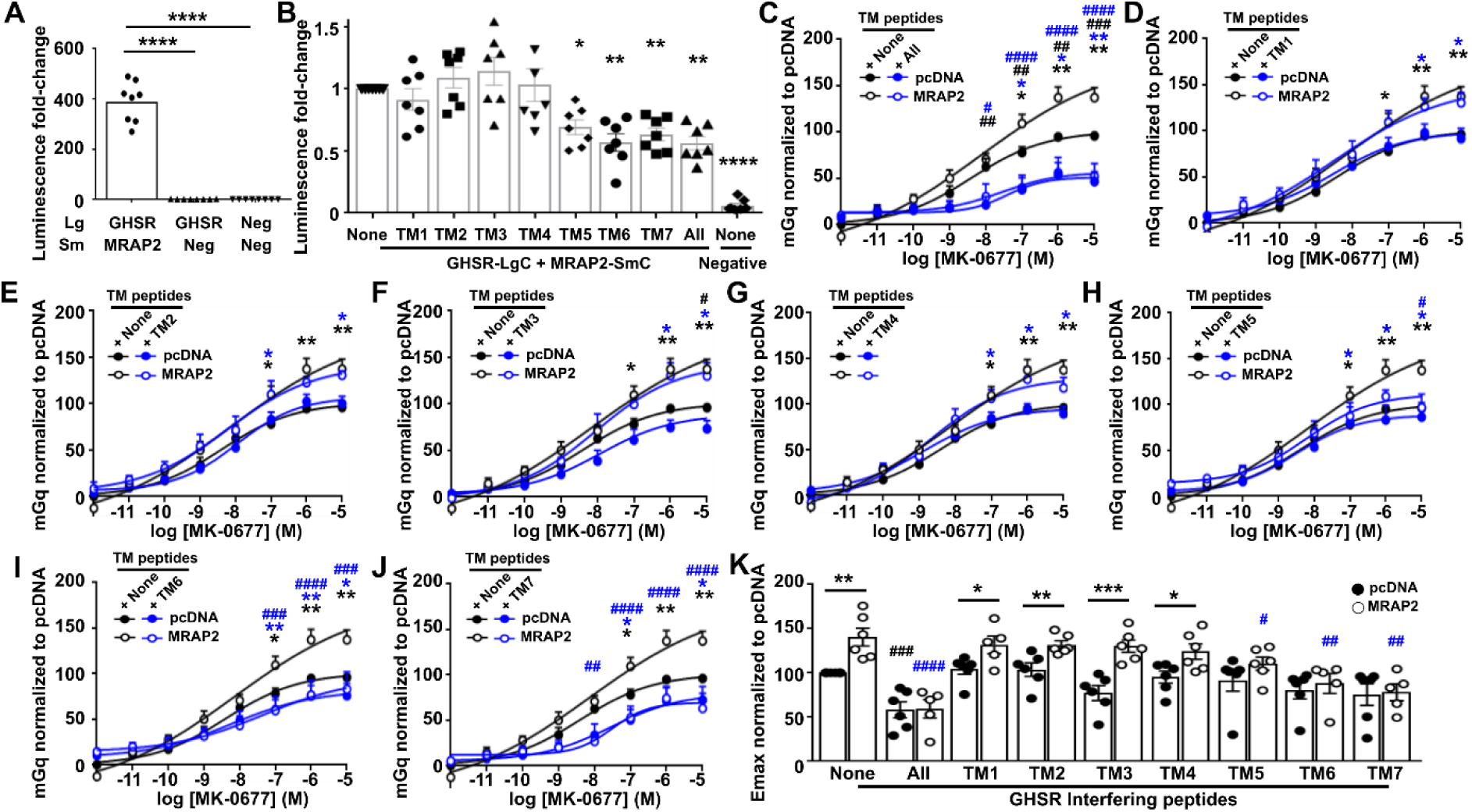
GHSR TM5-TM7 are important for GHSR-MRAP2 interactions and MRAP2-mediated signaling. (**A**) NanoBiT luminescence generated between GHSR-LgC and MRAP2-SmC, GHSR-LgC and SmC-empty. Values are normalized to those in cells transfected with LgC and SmC empty. N = 8. (**B**) NanoBiT luminescence generated between GHSR-LgC and MRAP2-SmC in cells exposed to GHSR TM interfering peptides. Values are normalized to those in cells exposed to vehicle. Statistical analyses were performed by one-way ANOVA with Sidak’s multiple-comparisons test compared to cells treated with vehicle. N = 7. (**C-J**) Normalized BRET responses between GHSR-Rluc8 and mGq in cells expressing MRAP2 or pcDNA and pre-incubated with (C) all TM peptides, or peptides targeting (D) TM1, (E) TM2, (F) TM3, (G) TM4, (H) TM5, (I) TM6, (J) TM7, with (K) Emax values. N = 5-6. Statistics for pEC50 are shown in Table S4. Statistical analyses in C-J were performed by two-way ANOVA with Tukey’s multiple comparisons test and compared: pcDNA none vs. MRAP2 none (black asterisk), pcDNA none vs. MRAP2 TM (blue asterisk), pcDNA none vs. pcDNA TM (black hash), MRAP2 none vs. MRAP2 TM (blue hash). Statistical analyses in K were performed by one-way ANOVA with Dunnett’s multiple-comparisons test and show comparisons between pcDNA none and pcDNA TM shown as a black hash and those between MRAP2 none and MRAP2 TM shown as a blue hash. ****p<0.0001, ***p<0.001, **p<0.01, *p<0.05.

To verify whether the TM interfering peptides affected MRAP2’s ability to facilitate signaling, BRET assays were performed to measure mGq recruitment to GHSR-Rluc8. Cells were transfected with mGq-Venus, GHSR-Rluc8 and pcDNA or MRAP2, and exposed to TM interfering peptides for two hours prior to measurement of BRET responses. Treatment of cells with all TM peptides simultaneously reduced signaling by both pcDNA and MRAP2-WT expressing cells, demonstrating that the peptides disrupt GHSR activity (Figure 4C). Exposure to peptides mimicking TM1-TM4 had no effect on MRAP2-mediated GHSR signaling, although the TM3-targeting peptide reduced signaling in cells transfected with pcDNA at the highest concentration of agonist (Figure 4D-G). Peptides targeting TM5, TM6 and TM7 reduced MRAP2-mediated increases in signaling and reduced the Emax, such that they were similar to pcDNA expressing cells (Figure 4H-K, Table S4). These peptides did not affect pcDNA responses, suggesting that the effect of peptides was on the MRAP2-GHSR interface or affected MRAP2’s ability to enhance GHSR signaling. AlphaFold2 did not predict a consistent interaction interface with MRAP2, although the model with the highest prediction score showed some interaction between MRAP2 and residues within TM5, TM6 and the TM7-ECL3 region (Figure S4). Therefore, it is likely that MRAP2 forms interactions with TM5, TM6 and TM7 of GHSR.

### The MRAP2 cytoplasmic region is critical for facilitating GPCR signaling

Our previous studies suggested the MRAP2 cytoplasmic region plays an important role in regulating GPCR signaling^8,17^. To investigate this further we designed three mutant proteins of MRAP2: one with truncation of the entire cytoplasmic region of MRAP2 (Truncation), the second with the extracellular and transmembrane region of MRAP1 with the MRAP2 cytoplasmic domain (N1C2), and a third with the MRAP2 extracellular and transmembrane region with the MRAP1 cytoplasmic domain (N2C1) (Figure 5A). The three mutant plasmids produced proteins with similar total protein expression (Figure 5B, S4A-C) and GPCRs were still expressed at the membrane (Figure S4D), although the cell surface expression of GHSR in the presence of the Truncation mutant was reduced compared to wild-type MRAP2 (Figure 5C). Deletion of the MRAP2 cytoplasmic region significantly reduced the ability of the accessory protein to facilitate GHSR-induced recruitment of mGq (Figure 5D, Table S5) and prevented the suppression in basal mGq responses compared to MRAP2-WT (Figure 5E). Additionally, GHSR-mediated elevations in Ca^2+^_i_ and reductions in cAMP were impaired by truncation of the MRAP2 cytoplasmic region (Figure 5F-G, Table S5). At high agonist concentrations, the Truncation mutant could increase signaling to significantly greater levels than the pcDNA control, indicating the extracellular and/or transmembrane region of GHSR also has a role in MRAP2-mediated signaling. Addition of the MRAP2 cytoplasmic region to the MRAP1 extracellular and transmembrane region, which is a similar length to MRAP2, was able to restore the ability of MRAP2 to enhance GHSR signaling, although replacement of the cytoplasmic region of MRAP2 with that from MRAP1 could not enhance GHSR-induced signaling above that observed for pcDNA (Figure 5D-G, Table S5). The Truncation mutant and N2C1 were similarly able to significantly reduce MC4R-and MC3R-mediated cAMP signaling when compared to MRAP2 WT, although loss of the cytoplasmic region impaired effects on constitutive activity (Figure 5H-J, Table S5). Thus, the MRAP2 cytoplasmic region has an important role in GPCR signaling.

**Figure 5.**
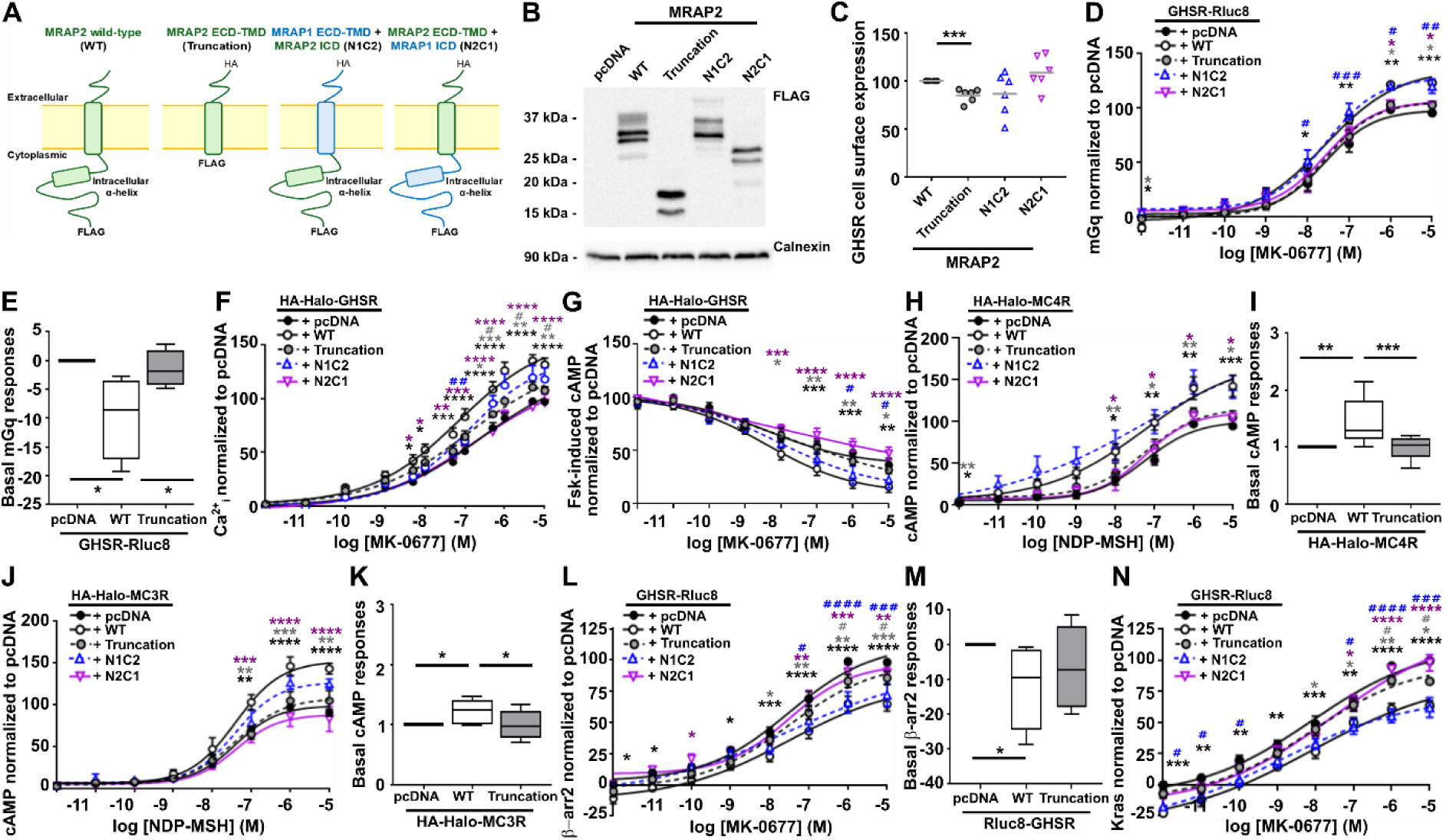
Loss of the MRAP2 cytoplasmic region impairs MRAP2’s ability to facilitate GPCR signaling. (**A**) Cartoon showing the full-length WT MRAP2 protein, the MRAP2 Truncation protein and the two MRAP2 and MRAP1 chimeric proteins. (**B**) Western blot analysis of MRAP2-FLAG mutant constructs. Densitometry from four blots, with full blots shown in Figure S4. (**C**) Cell surface expression of GHSR-Rluc8 in the presence of the MRAP2 mutant proteins. Gray line shows mean. N=6. Cells were transfected with either pcDNA, MRAP2 WT or MRAP2 mutant proteins and the following assessed: (**D**) Normalized BRET responses between GHSR-Rluc8 and mGq, with (**E**) constitutive activity (N=4),(**F**) GHSR-mediated intracellular calcium (Ca^2+^_i_) responses measured by Fluo-4 (N=8), (**G**) Suppression of forskolin-induced cAMP responses in cells transfected with HA-Halo-GHSR (N = 5). (**H-I**) cAMP responses to NDP-MSH in cells transfected with (H) HA-Halo-MC4R with (I) basal activity or (J) HA-Halo-MC3R with (K) constitutive activity (N = 8). (**L**) Normalized BRET responses with (**M**) constitutive activity between GHSR-Rluc8 and β-arrestin-2-YFP (N=7) and (**N**) normalized BRET responses between GHSR-Rluc8 and Venus-Kras (N=6). Responses are expressed relative to the pcDNA maximum set at 100. Data shows mean±SEM in D, F-H. J, L, N and mean with maximum and minimum in E, I, K, M. Statistical analyses were performed by one-way ANOVA with Dunnett’s multiple-comparisons test in C, E, I, K, M and two-way ANOVA with Sidak’s multiple-comparisons test in other panels. Comparisons in D, F-H, J, L, N show: WT vs. pcDNA (black asterisk), WT vs. Truncation (gray asterisk), WT vs. N1C1 (purple asterisk), pcDNA vs. Truncation (gray hash), pcDNA vs. N1C2 (blue hash). pEC50 values are shown in Table S5. ****p<0.0001, ***p<0.001, **p<0.01, *p<0.05.

The cytoplasmic region of MRAP2 may enhance GPCR signaling, at least in part, by impairing the ability of β-arrestin to couple to membrane bound receptors, thus prolonging the duration of active receptors at cell surfaces. Consistent with this, deletion of the MRAP2 cytoplasmic region or replacement with the MRAP1 cytoplasmic region enhanced β-arrestin-2 recruitment to GHSR without affecting the basal response (Figure 5L-M, Table S5), and enhanced internalization from plasma membranes expressing Kras (Figure 5N, Table S5). To verify these findings, colocalization of HA-SNAP-GHSR with Venus-Kras was monitored in cells prior to, and 30 minutes following, exposure to agonist. GHSR and Kras were expressed at the cell surface under basal conditions in the presence of pcDNA, MRAP2-WT or MRAP2-Truncation plasmids (Figure 6A). Following exposure to agonist, GHSR was observed in intracellular vesicles, and colocalization with the plasma membrane marker Kras was reduced. Internalization was impaired in cells expressing MRAP2-WT compared to those expressing pcDNA and the MRAP2-Truncation (Figure 6A-C). GHSR was also recruited to Rab5-positive endosomes more rapidly in the presence of pcDNA or the MRAP2-Truncation plasmid following agonist exposure (Figure 6D-F).

**Figure 6.**
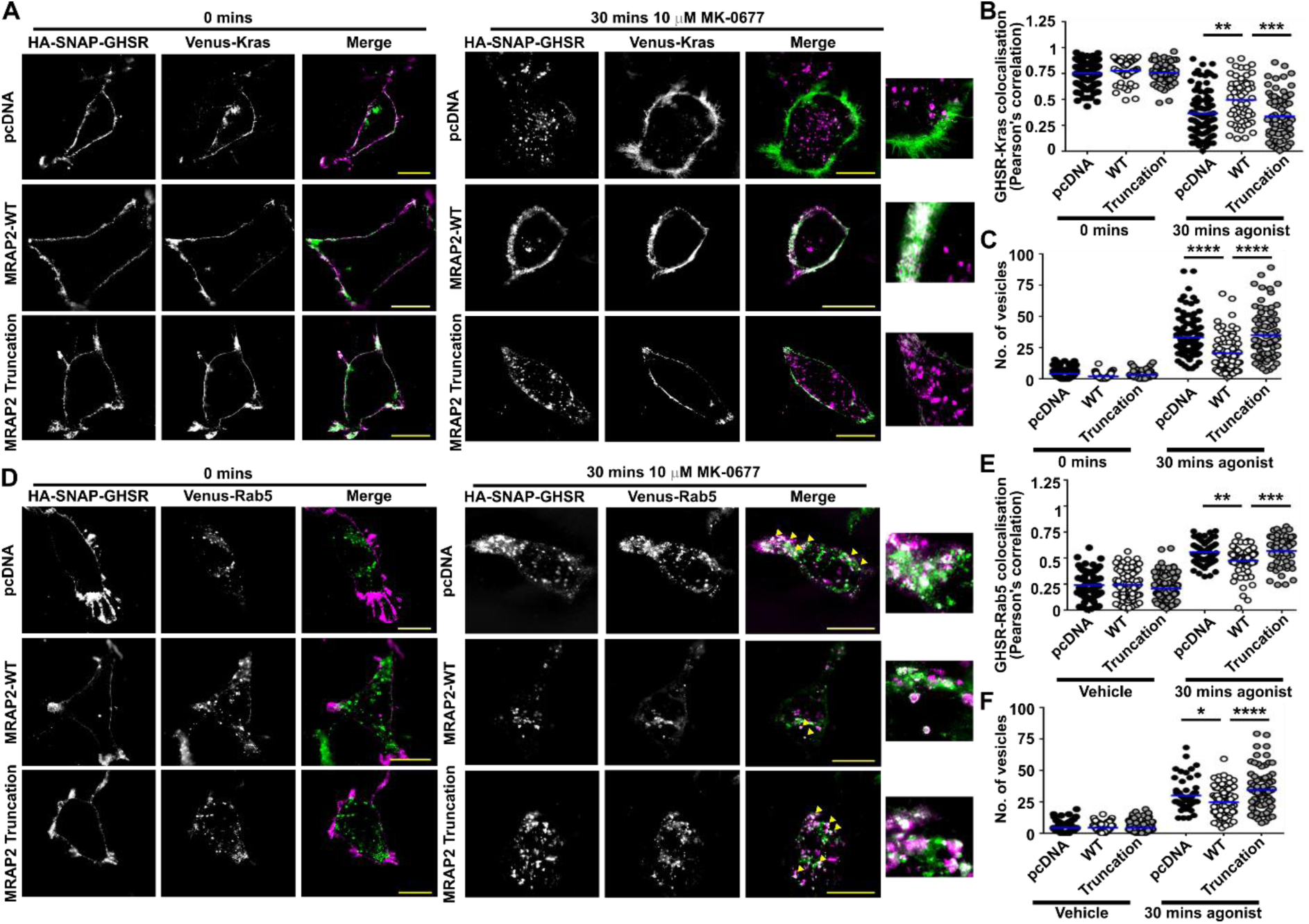
Loss of the MRAP2 cytoplasmic region enhances GPCR internalization. (**A**) SIM imaging of GHSR and Venus-Kras with pcDNA, MRAP2-WT or the MRAP2 Truncation mutant with (**B**) correlation between GHSR and Kras assessed by Pearson’s coefficient, and (**C**) quantification of the number of internalized vesicles. Number of cells from four independent transfections for each group as follows: pcDNA vehicle (118) and agonist (106), MRAP2-WT vehicle (63) and agonist (75), MRAP2-Truncation vehicle (61) and agonist (71). **(D**) SIM imaging of GHSR and Venus-Rab5 with pcDNA, MRAP2-WT or the MRAP2 Truncation mutant, with (**E**) correlation between GHSR and Rab5 assessed by Pearson’s coefficient, and (**F**) Quantification of the number of internalized vesicles. Number of cells from 4-5 independent transfections for each group as follows: pcDNA vehicle (67) and agonist (56), MRAP2-WT vehicle (95) and agonist (80), MRAP2-Truncation vehicle (120) and agonist (77). Yellow arrows show areas of colocalization in A and D. Blue line shows mean in B, C, E, F. Scale, 5 μm. Statistical analyses were performed by one-way ANOVA with Dunnett’s multiple-comparisons test in B, C, E, F. ****p<0.0001, ***p<0.001, **p<0.01, *p<0.05.

### Obesity-associated variants in the MRAP2 cytoplasmic region enhance constitutive activity

As our studies of the MRAP2 cytoplasmic domain demonstrated its importance in GPCR signaling we hypothesized that MRAP2 variants associated with human obesity in this region would affect GHSR signaling. Five MRAP2 variants within the cytoplasmic region that have been identified in individuals with overweight and/or obesity were tested alongside two variants (V91A, H133Y) that have only been identified in normal weight individuals. GHSR was expressed at the cell surface in the presence of all the MRAP2 variants (Figure S6). Recruitment of mGq to GHSR in the ligand-free state was enhanced by four of the cytoplasmic region variants (R113G, S114A, L115V, N121S), with significantly reduced maximal response to agonist in cells expressing R113G (Figure 7A-B, Figure S7A, Table S6). Four MRAP2 variants (R113G, S114A, L115V, T193A) associated with overweight and/or obesity also enhanced constitutive IP3 generation and two impaired maximal responses to ligand (Figure 7C-D, Figure S7B, Table 6). All five obesity-associated variants enhanced basal intracellular calcium responses and reduced maximal responses (Figure 7E-F, Figure S7C, Table S6). The ability of GHSR to suppress forskolin-induced cAMP was also impaired by all five MRAP2 variants (Figure 7G, Figure S8, Table S6). Thus, human variants in the cytoplasmic region that are associated with human obesity enhance GHSR constitutive activity.

**Figure 7.**
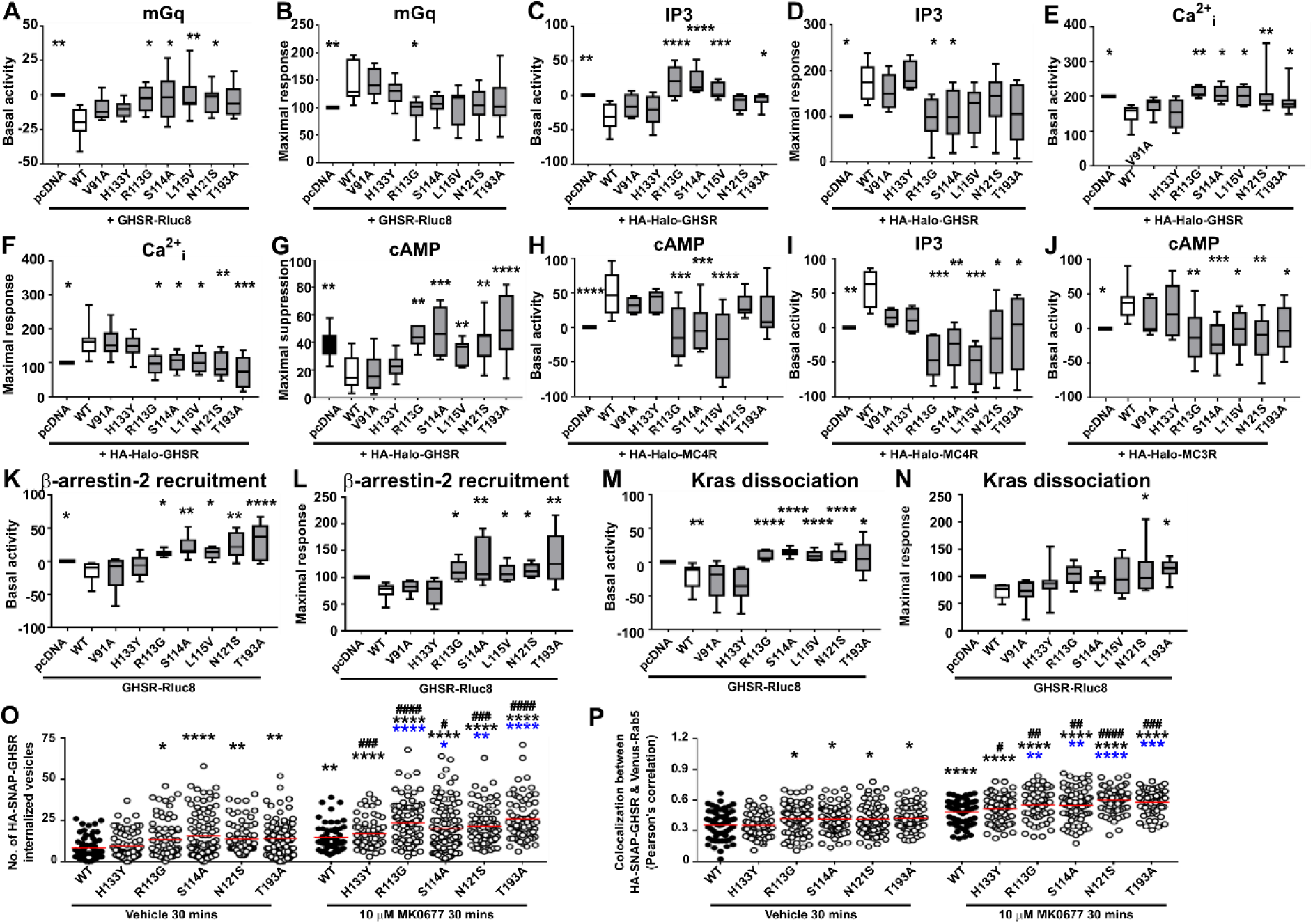
Effect of MRAP2 human obesity-associated variants on GHSR signaling. Cells were transfected with either pcDNA, MRAP2 WT or MRAP2 mutant proteins and the following assessed: (**A**) GHSR-mediated basal activity and (**B**) maximal responses from BRET between GHSR-Rluc8 and Venus-mGq (N = 6-8), (**C**) GHSR-mediated basal activity and (**D**) maximal responses from IP3 NanoBiT biosensor assays (N = 6-7), (**E**) GHSR-mediated basal activity and (**F**) maximal responses from intracellular calcium (Ca^2+^_i_) measured by Fluo-4, (**G**) GHSR-mediated suppression of forskolin-induced increases in cAMP (N = 7-10). (**H-I**) MC4R-mediated basal activity from (H) cAMP (N = 5-7) and (I) IP3 (N = 4-6). (**J**) MC3R-mediated basal cAMP activity (N = 5-7). (**K**) Basal activity and (**L**) maximal responses between GHSR-Rluc8 and β-arrestin-2-YFP (N = 5-7). (**M**) Basal activity and (**N**) maximal responses from BRET between GHSR-Rluc8 and Venus-Kras (N = 7-8). Data shows minimum and maximum in all panels. Concentration-response curves for mGq BRET, IP3 and Fluo-4 assays in Figure S7, for cAMP in Figure S8, and for β-arrestin-2 and Kras in Figure S9. pEC50 values are shown in Table S6. Full dose-response curves for H-J have been published^8,17^. Statistical analyses were performed by one-way ANOVA with Dunnett’s multiple-comparisons test. Maximal responses at each concentration were measured for Fluo-4 and responses expressed relative to the pcDNA maximal response set at 100. For cAMP, responses were normalized to the basal response for each group. For other assays AUC was measured and responses expressed relative to the pcDNA maximal response set at 100. (**F**) Quantification of the number of internalized vesicles and (**G**) correlation between GHSR and Venus-Rab5 assessed by Pearson’s coefficient in SIM images of cells exposed to vehicle or MK06-77 for 30 minutes. Images are shown in Figure S10. Statistical analyses were performed by one-way ANOVA with Dunnett’s multiple-comparisons test and compare responses to WT vehicle (black asterisk), WT agonist (blue asterisk) or vehicle vs. agonist within each variant (black hash). ****p<0.0001, **p<0.01, *p<0.05.

As we have shown that MRAP2 increases MC4R and MC3R constitutive activity (Figure 5), we hypothesized that MRAP2 variants may also affect constitutive activity of these receptors. We reanalysed our previous datasets^8,17^ and found that MRAP2 variants reduce constitutive activity of MC4R and MC3R (Figure 7H-J). Therefore, MRAP2 variants have a significant effect on constitutive activity of multiple GPCRs.

### MRAP2 cytoplasmic variants enhance Barr2 recruitment and internalization

We then assessed the effects of the MRAP2 obesity-associated variants on GHSR internalization. All five MRAP2 variants enhanced β-arrestin recruitment by increasing the basal activity and enhancing maximal responses (Figure 7K-L, Figure S9A, Table S6). There was a significant reduction in pEC50 for cells transfected with the five MRAP2 variants and pcDNA when compared to MRAP2-WT (Table S6). Dissociation of GHSR from the plasma membrane marker Kras was also impaired, with five variants exhibiting enhanced basal activity, and two variants (N121S, T193A) enhancing maximal responses (Figure 7M-N, Figure S9B, Table S6). The V91A and H133Y variants which have been described in individuals with normal weight did not impair GHSR internalization. Trafficking of GHSR was further assessed by SIM with HA-SNAP-GHSR labelled with anti-HA antibody and colocalization with Rab5 imaged following a 30-minute exposure to either vehicle or 10 µM MK0677 (Figure S10A). Four obesity-associated and one variant identified in normal weight individuals were investigated. The number of internalized vesicles was higher in cells expressing the obesity-associated variants following exposure to both vehicle and MK0677 (Figure 7O, Figure S10A-B). Similarly, colocalization between GHSR and Rab5, measured by Pearson’s coefficients, were increased in cells expressing the four obesity-associated GHSR variants, while cells expressing H133Y had similar colocalization to MRAP2-WT expressing cells (Figure 7P, Figure S10A-B). To determine whether there was a significant difference between agonist-driven responses in each group we assessed the fold-change increase between constitutive internalization and agonist-driven internalization in each data set. This showed that there was no significant difference in fold-change response between vehicle and MK0677, except for S114A in one dataset (Figure S10C-D). Therefore, obesity-associated variants increase GHSR internalization by enhancing basal β-arrestin recruitment and constitutive internalization.

## Discussion

Our studies reveal new insights into the shared mechanisms by which MRAP2 affects GPCR activity. These include enhanced signaling by multiple signaling pathways, impaired recruitment of β-arrestin-2 binding and reduced receptor internalization that likely prolongs the duration of active receptor at cell surfaces contributing to increased signaling. Single-molecule studies of MRAP2 in complex with either GHSR or MC4R revealed that heterodimers are preferentially formed between monomeric MRAP2 and receptor. This is consistent with our previous analyses of MC3R and MRAP2 heterodimers that similarly formed 1:1 complexes^8^, and with models of GPCRs with MRAP2^8,17,18,39^. This provides further evidence that in vitro studies of GPCRs with MRAP2 should be conducted with equal concentrations of the two proteins rather than with saturating concentrations of MRAP2 that could lead to overexpression artefacts and are inconsistent with expression levels in neuronal cells^8^. In contrast to previous studies^2,16^ we were able to demonstrate that expression of equal DNA concentrations of GHSR and MRAP2 had robust effects on signaling. While there were differences in cell-lines used (CHO in previous studies and HEK in the current studies) we were also able to replicate our findings in mouse hypothalamic cell-lines (mHypoN-39), which are more reflective of the neuronal physiological environment. Therefore, we propose that future studies should examine MRAP2 effects on GPCRs in cells transfected with equal protein concentrations.

GHSR and MC4R were more likely than MC3R to form heterodimers comprising either monomeric GPCR with dimeric MRAP2 or complexes with two photobleaching steps for GPCR and MRAP2. MRAP2 has been shown to form dimers^2^ and our previous SiMPull analyses showed MRAP2 dimers are present at cell membranes, although monomeric MRAP2 is more common^8^. It is possible that MRAP2 oligomerises at cell membranes under basal conditions until GPCRs are present, with which it forms stable complexes. In agreement with previous studies^40,41^ we found that GHSR and MC4R form dimers and higher-order oligomers at cell membranes when expressed alone, and found that co-transfection with MRAP2 disrupted oligomerization of all three GPCRs. The role of MC4R oligomerization is incompletely understood. MC4R mutations that disrupt oligomerization are pathogenic^42^. However, this may be due to dominant-negative effects of the mutant GPCR on the wild-type receptor, which was reported in earlier studies^41^. Later studies indicated that MC4R couples to Gs more effectively following dimer separation, and that MRAP2 potentiates signaling by promoting disruption of GPCR oligomerization^18,43^. Whether GHSR oligomerization engenders biased signaling as demonstrated for some GPCRs^44^, or impairs G protein coupling is not known. Our studies show that MRAP2 disrupts oligomerization of all three GPCRs investigated, suggesting that this may be a canonical mechanism employed by MRAP2 and GPCRs to form heterodimers.

The formation of complexes between GHSR and MRAP2 most likely involves residues within TM5, TM6 and TM7 of GHSR. Cell-penetrating transmembrane interfering peptides mimicking these helices disrupted complex formation and recruitment of mGq protein. Structural studies have shown agonist engagement with the bifurcated ligand-binding pocket of GHSR conveys conformational changes within an aromatic cluster at TM6-TM7 and shifts TM7 toward TM6 to activate the receptor^37,45^. MRAP2 may act as an allosteric modulator to promote adoption of this conformation more readily upon ligand binding. Modeling and mutagenesis studies indicate that MRAP2 similarly engages with MC4R and MC3R TM5 and TM6 and we have previously proposed that MRAP2 interaction more readily allows formation of the large cytoplasmic cavity between TM5-TM7 that accommodates G protein binding to melanocortin receptors^8,46,47^. Our findings that GHSR binding to MRAP2 similarly involves these transmembrane helices suggests this is a shared mechanism by which MRAP2 binds to GPCRs.

GHSR has considerable basal activity^20^ and IP3 ligand-independent signalling has been shown to reduce when MRAP2 is overexpressed^16^. We showed that MRAP2 retains its ability to reduce GHSR constitutive activity when the two proteins are expressed at equal concentrations. Reduced basal activity could allow the receptor a larger dynamic range over which agonist could activate the receptor. However, the A204E mutant that lacks constitutive activity is regarded as a loss-of-function mutation as its physiological responses (increased food intake, growth hormone release) are lost^24,25^, despite the receptor retaining its ability to increase signalling in the presence of ghrelin. Therefore, MRAP2 binding to GHSR must also facilitate a conformation that allows the receptor to more readily respond to ligand. Studies of GHSR in lipid nanodiscs have suggested that GHSR may exist in multiple ligand-free states in which the preassembled GHSR-Gq complex shuttles between inactive and active states, and that ligand increases the population of active states ^40^. It is therefore possible that MRAP2 binding to GHSR favours a conformation that bypasses the need for these intermediate states and readily adopts the ligand-bound active state. Our studies show that MRAP2 likely interacts with TM6 and TM7 that are known to undergo large conformational changes upon ligand binding to open the G protein binding cavity^48^, and it is possible that MRAP2 binding favours this open conformation. More recent studies suggest the lipid environment, particularly the presence of PIP2, holds G proteins in proximity to GHSR, and this enables rapid formation of a stable complex with G proteins^49^. It is possible that MRAP2 binding to GHSR alters the lipid environment to increase PIP2 and G proteins located close to GHSR. Future studies investigating how the lipid environment affects MRAP2 or structures of GHSR in complex with MRAP2 will help elucidate these mechanisms.

Deletion of the MRAP2 cytoplasmic region reduced its ability to enhance G protein signaling and its effect on constitutive activity. This suggests that the cytoplasmic region is important for GPCR activity and possible mechanisms could include helping the receptor adopt its active conformation, affecting the lipid composition of the membrane to favour PIP2 and Gq proximity^49,50^ or occluding the β-arrestin binding site. Previous studies have shown that MRAP2 blocks GRK2 and PKC access to GHSR intracellular loops that are required for the phosphorylation events that precede β-arrestin recruitment^51^, and our studies show deletion of the MRAP2 cytoplasmic domain increases β-arrestin recruitment to GHSR, consistent with a role for the cytoplasmic region of MRAP2 in this blocking event. In agreement with this, MRAP2 impairs GPCR constitutive and agonist-driven internalization, while loss of the MRAP2 cytoplasmic region, which would allow phosphorylation and β-arrestin-2 recruitment to go unhindered, enhances internalization. Collectively, these events prolong the presence of active receptor at cell surfaces, such that overall signaling is enhanced. However, MRAP2 retains some ability to enhance GPCR signaling in β-arrestin-depleted cells^16^, suggesting the mechanism cannot solely depend on impaired β-arrestin recruitment. Ubiquitination is also known to promote GHSR and MC4R internalization^39,52^ and removal of the MRAP2 cytoplasmic region may also allow ubiquitination to occur more readily. Our studies also showed that despite structural similarities, the MRAP1 cytoplasmic domain cannot compensate for the MRAP2 domain, although the transmembrane regions may have similar modes of binding or effects on receptor activity.

The overweight and obesity phenotypes observed in individuals with MRAP2 variants can be explained by the reduced MC4R activity associated with these mutant proteins^17,29,43^. However, additional conditions (hyperglycaemia, hypertension)^29^ reported in individuals with MRAP2 variants suggest other GPCRs may be affected, and we examined the effect of variants in the MRAP2 cytoplasmic region on GHSR signaling and trafficking. The five variants had variable effects on receptor maximal responses, although variants consistently enhanced GHSR constitutive activity by all signaling pathways. Constitutive activity regulates multiple GHSR physiological functions. In mice with an A203E mutation that ablates constitutive activity, food intake and energy homeostasis are impaired and blood glucose is reduced^25^, and there is some evidence that humans with variants in GHSR that impair constitutive activity have lower BMI in early life^32,33^. Therefore, the increased constitutive activity observed with the MRAP2 variants would be expected to increase food intake and increase blood glucose, which could contribute to some of the additional conditions observed in individuals with MRAP2 variants. Effects on growth are also observed in mice and humans with impaired GHSR constitutive activity^24,25,30^, but effects on height have not been reported in individuals with MRAP2 variants. However, linear growth is a highly complex trait with over 100 loci associated with human height^53^ and GHSR and MRAP2 will only partially contribute to this. Moreover, as we have shown that LEAP2 is still able to reduce GHSR activity in the presence of MRAP2, and previous studies have shown LEAP2 can reduce GHSR constitutive activity^12,13^, symptoms that arise in patients with MRAP2 variants will likely depend on a balance between effects on GHSR, LEAP2, and other GPCRs. This balance could also explain why deletion of *Mrap2* on different genetic backgrounds had different effects on insulin and glucose handling^15,28^.

Although there are shared mechanisms by which MRAP2 interacts with and modulates GPCR function we did detect some differences. In contrast to GHSR in which basal activity was reduced, we identified an increase in MC4R and MC3R constitutive activity. While MC4R has been reported to harbor some basal activity, MC3R has minimal constitutive activity^54^, and therefore reductions in constitutive activity would not increase the dynamic range of ligand responses in the same way as GHSR that has high constitutive activity. Instead, constitutive activity for melanocortin receptors is understood to provide a tonic satiety signal for maintaining long-term energy homeostasis^55^. Enhancement of this signal by MRAP2 would be consistent with its known role in enhancing MC4R function. Obesity-associated MRAP2 variants reduced MC4R and MC3R constitutive activity similarly to truncation of the MRAP2 cytoplasmic region, and this reduction in the tonic satiety signal could contribute to the weight gain observed in individuals with these genetic variants. It is also possible that oligomerization of MC3R and MC4R prevents high constitutive activity, as it does agonist-driven signaling, as they may be less able to couple to G proteins or other signal components.

In summary, our studies have identified critical mechanisms by which MRAP2 regulates GPCR signaling and trafficking. We have provided further evidence that MRAP2 has an essential role in energy homeostasis and that MRAP2 genetic variants may modulate signaling by multiple GPCRs to cause overweight and obesity.

## Materials & Methods

### Plasmid constructs and compounds

A full list of plasmids with their source can be found in Table S1. For single molecule pull-down experiments, constructs were generated with an N-terminal signal peptide from rat mGluR2^34^, followed by affinity tags (HA or FLAG), self-labeling protein tags capable of conjugation to organic dyes (SNAP, CLIP, or Halo), and human GHSR, MC4R or MC3R and MRAP2. Cloning into the pRK5 vector was performed using reagents from Promega and oligonucleotides from Sigma to generate ss-HA-Halo, ss-HA-SNAP and ss-FLAG-CLIP plasmids (Table S1). For the MRAP2 truncation plasmid, MRAP2 amino acids 1-63 were cloned into pcDNA3.1 with an N-terminal HA tag and a C-terminal FLAG tag. For the N1C2 plasmid, MRAP1 residues 1-56 were cloned with an N-terminal HA tag, followed by MRAP2 amino acids 64-205 and a C-terminal 3xFLAG tag. For MRAP2-N1C2, MRAP2 residues 1-63 were cloned following an HA tag, with MRAP1 amino acids 57-172 and a 3xFLAG tag. All plasmids were sequence-verified by Source Bioscience. MK0677 (Tocris), NDP-MSH (Cambridge Bioscience) and LEAP2 (MedChemExpress) were used at the concentration stated in each figure. Transmembrane interfering peptides were custom synthesized by Genscript with a purity of ≥95%. Peptides were designed with an intracellular TAT cell-penetrating sequence and comprised the following sequences: PAPLLAGVTATCVALFVVGIAGNLLTMLVVSRFGRKKRRQRRRPPQGG (TM1), GRKKRRQRRRPPQGGTTNLYLSSMAFSDLLIFLCMPLDLVRLWQ (TM2), LTITALSVERYFAICGRKKRRQRRRPPQGG (TM3), GRKKRRQRRRPPQGGTKGRVKCSA (TM4), GLLTVMVWVSSIFFFLPVFCLTVLYSLIGRKLWGRKKRRQRRRPPQGG (TM5), GRKKRRQRRRPPQGGRDQNHKQTVKMGRYLFSKSF (TM6), QYCNLVSFVLFYLSAAINPILYNIMSGRKKRRQRRRPPQGG (TM7). Peptides were diluted in water to a 1mM stock, then used at 10 µM in assays.

### Cell culture and transfection

Adherent HEK293 (AdHEK) cells (Agilent Technologies, AD-293, RRID:CVCL_9804) and mouse hypothalamic cells mHypoE-39 (Cedarlane Cellutions Biosystems Inc. (mHypo-N39, RRID:CVCL_D439)) were maintained in DMEM-Glutamax media (Merck) with 10% calf serum (Invitrogen) at 37°C, 5% CO_2_. Cells were routinely screened to ensure they were mycoplasma-free using the TransDetect Luciferase Mycoplasma Detection kit (Generon). Expression constructs were transiently transfected using Lipofectamine 2000 (LifeTechnologies), following manufacturer’s instructions.

### Western blot analysis

For MRAP2 mutant expression studies, cells were transfected with 1 µg per well in a 6-well plate of either pcDNA, MRAP2-WT, MRAP2-Truncation or MRAP2-MRAP1 chimeric mutants. Cells were lysed 48-hours later in NP40 buffer and western blot analysis performed. Blots were blocked in 5% marvel/TBS-T, then probed with 1:1000 mouse anti-FLAG (M2 antibody, Sigma-Aldrich, RRID:AB_262044) and 1:1000 rabbit anti-calnexin (Millipore, Cat# AB2301, RRID:AB_10948000) or 1:1000 rabbit alpha-tubulin (Abcam, Cat# ab176560, RRID:AB_2860019) antibodies. Blots were visualized using the Immuno-Star WesternC kit (BioRad) on a BioRad Chemidoc XRS+ system. Densitometry was performed using ImageJ (NIH), and protein quantities normalized to calnexin or alpha-tubulin housekeeper.

### NanoBiT interaction assays

GHSR was cloned into the LgBiT-C plasmid (Promega). The MRAP2-SmC construct has been described^8^. HEK293 cells were seeded at 10,000 cells/well in 96-well plates and transfected the same day with 100ng of GHSR-LgC and SmC-MRAP2, GHSR-LgC and LgC-empty, or LgC-empty and SmC-empty as a negative control (Promega). Following 48-hours, media was changed to FluoroBrite DMEM phenol red-free media (Gibco) with 10% calf serum (FluoroBrite complete media) with 40 μL Nano-Glo substrate (Promega) and luminescence signals read (duplicate for each condition) on a Glomax (Promega) plate reader at 37°C. Data was normalized to luminescence values in the negative control for interaction assays. For studies with TM interfering peptides, cells were pre-incubated with 10 µM of each peptide, or an equal volume of water for the vehicle, for 2 hours. Luminescence signals were then read (duplicates for each condition) and data was normalized to luminescence values in wells transfected with GHSR-LgC and SmC-MRAP2 that were treated with vehicle.

### Bioluminescence resonance energy transfer (BRET) assays

NanoBRET assays were performed using methods adapted from previous studies^56^. AdHEK or mHypo-N39 cells were seeded at 10,000 cells/well in 96-well white plates and transfected the same day with 50ng GHSR-Rluc8 and 500ng Venus-mGq or Venus-β-arrestin-2 or Venus-Kras and 50ng of either pcDNA3.1 or 3xFLAG-MRAP2. Forty-eight hours later, cells were washed with PBS, then 40μl of 10μM coelenterazine-H (Promega) diluted in FluoroBrite complete media were added to the wells. BRET measurements were recorded using a GloMax microplate reader (Promega) at donor wavelength 475-30 and acceptor wavelength 535-30 at 37°C. The BRET ratio (acceptor/donor) was calculated for each time point. Four baseline recordings were made, then agonist added at 8 minutes and recordings made for a further ∼40 minutes. The average baseline value recorded prior to agonist stimulation was subtracted from the experimental BRET signal. All responses were then normalized to that treated with vehicle to obtain a normalized BRET ratio. AUC was calculated in GraphPad Prism and these values used to plot concentration-response curves with a 4-parameter sigmoidal fit. For assessment of cell surface expression, BRET values prior to agonist stimulation were compared in cells transfected with GHSR-Rluc8, Venus-Kras and either pcDNA, MRAP2-WT or MRAP2-variants. For studies with TM interfering peptides, cells were pre-incubated with 10 µM of each peptide, or an equal volume of water for the vehicle, for 2 hours. For studies with LEAP2, cells were pre-incubated with 1, 10 or 100 nM for 30 minutes prior to performance of the assays.

### Fluo-4 intracellular calcium assays

AdHEK cells were plated in black sided 96-well plates and transfected with 50ng HA-SNAP-GHSR or HA-Halo-GHSR and either 50ng pcDNA3.1 or 3xFLAG-MRAP2 (WT or mutant) plasmids. Cells were incubated for 48 hours prior to performance of the Fluo-4 intracellular calcium assays. Fluo-4 AM (Molecular Probes, Life Technologies) was resuspended in DMSO/ 0.03% pluronic acid, then a working solution made on the day of the experiment at 1:1000 dilution in HBSS (Sigma) with 10mM HEPES (Gibco) (HANKS-HEPES solution). Cells were incubated with 50µL Fluo-4 working solution and incubated at room temperature for one hour. Cells were washed once in HANKS-HEPES solution, then 50µL fresh HANKS-HEPES added prior to assay performance on a Glomax Discover at 37°C with excitation 494 nm and emission at 516 nm. Ten baseline reads were made per well prior to injection of drugs from preloaded injectors using an automated protocol. Following injection of each drug concentration, ten reads were performed. At least three technical replicates were performed per condition and the position of each condition varied across each plate. Data was plotted in GraphPad Prism. The maximal response for each drug concentration was derived, normalized to baseline responses of the MRAP2-WT transfected wells and concentration-response curves plotted with a 4-parameter sigmoidal fit.

### Single molecule pull-down (SiMPull)

Cells were seeded in 12-well plates and transfected with 300 ng of Halo-tagged plasmids and 600ng of CLIP-tagged plasmids. After 24 hours, cells were washed with extracellular solution (comprising 135 mM NaCl (Sigma), 5.4 mM KCl (Sigma), 10 mM HEPES (Gibco), 2 mM CaCl_2_ (VWR Chemicals); 1 mM MgCl_2_ (Sigma), pH 7.4), then labelled with 2 μM of cell-membrane impermeable dyes (CLIP-surface 547 (BC-DY547, NEB) for FLAG-CLIP tagged plasmids, or CA-Sulfo646 ^35^ for HA-Halo tagged plasmids) in extracellular solution for 45 min at 37 °C. Cells were washed with extracellular solution, harvested in 1x Ca^2+^-and Mg^2+^-free PBS, then cell pellets lysed in buffer (Tris pH8, NaCl, EDTA (all from Sigma)) containing 0.5% Lauryl Maltose Neopentyl Glycol/ 0.05% Cholesteryl Hemisuccinate (LMNG-CHS) (Anatrace) and protease inhibitor (Roche). Microflow chambers were prepared by passivating a glass coverslip and quartz slide with mPEG-SVA and biotinylated PEG (MW = 5000, 50:1 molar ratio, Laysan Bio), as previously described^34,57^. Prior to each experiment a chamber was incubated with 0.2 mg/ml NeutrAvidin (Fisher Scientific UK) for 2 min, washed in T50 buffer (50 mM NaCl, 10 mM Tris), then incubated with 10 nM biotinylated anti-HA antibody (ab26228, Abcam, RRID:AB_449023) in T50 buffer (50 mM NaCl, 10 mM Tris) for 30 minutes. Fresh cell lysates were mixed with dilution buffer (1:10 lysis working solution with extracellular solution) and added to the flow chamber until a suitable single molecule spot density was obtained. Chambers were washed with dilution buffer to remove unbound receptor, then single molecule movies obtained as previously described ^34^ using a 100x objective (NA 1.49) on an inverted microscope (Olympus IX83) with total internal reflection (TIR) mode at 20 Hz with 50 ms exposure time with two sCMOS cameras (Hamamatsu ORCA-Flash4v3.0). Samples were excited with 561 nm and 640 nm lasers to excite BC-DY547 and CA-Sulfo-646, respectively. Single molecule movies were recorded sequentially at 640 nm, then 561 nm until most molecules were bleached in the field. Images were analyzed using a custom-built LabVIEW program ^58^. Each movie was concatenated using MatLab (R2022a), then loaded on LabVIEW to visualize each channel for co-localized molecules. The fluorescence trace of each molecule was inspected manually and bleaching steps aligned. Data were plotted using GraphPad Prism.

### Structured illuminated microscopy (SIM)

Cells were plated on 24 mm coverslips (VWR) and transfected with 500ng of each plasmid 36-hours prior to experiments. For studies of cell surface expression, cells were fixed with 4% PFA in PBS and exposed to 1:1000 anti-HA mouse monoclonal antibody (BioLegend Cat#901514, RRID:AB_2565336) or 1:1000 anti-FLAG mouse monoclonal antibody (M2, Sigma-Aldrich), followed by Alexa Fluor 647 goat anti-mouse (Cell Signaling Technology Cat# 4410, RRID:AB_1904023). For FLAG-CLIP-GHSR and HA-SNAP-MRAP2 colocalization studies, the anti-HA rabbit primary antibody (ab26228, Abcam) was used with the anti-FLAG antibody, followed by Alexa Fluor 647 goat anti-mouse and Alexa Fluor 488 goat anti-rabbit (Cell Signaling Technology Cat# 4412, RRID:AB_1904025). For studies with Venus-Kras and HA-SNAP-GHSR or Venus-Rab5 and HA-SNAP-GHSR, cells were exposed to 1:1000 anti-HA mouse monoclonal antibody (BioLegend Cat#901514, RRID:AB_2565336) with either vehicle or 10 μM MK-0677 for 30 minutes. Cells were fixed, permeabilized and exposed to the Alexa Fluor 647 secondary antibody. Samples were imaged on a Nikon N-SIM system (Ti-2 stand, Cairn TwinCam with 2 × Hamamatsu Flash 4 sCMOS cameras, Nikon laser bed 488 and 647 nm excitation lasers, Nikon 100 × 1.49 NA TIRF Apo oil objective). SIM data was reconstructed using NIS-Elements (v. 5.21.03) slice reconstruction. Colocalization and Pearson’s correlation coefficient was measured using the ImageJ plugin JACoP. Vesicles were counted in ImageJ using protocols previously described^59^. A manual threshold was set for each image to subtract background. Images were then processed using ‘Process>Binary>Convert to mask’, then ‘Analyze particles’ in ImageJ. Images were manually inspected to ensure vesicles were captured by this method and observers were blinded to the conditions of the experiment.

### IP_3_ biosensor

IP3 assays were performed as described^60^. AdHEK cells were plated in white 96-well plates and transfected the same day with 200ng LgBiT-IP_3_R2-SmBiT plasmid, 50ng HA-SNAP-GHSR or HA-Halo-GHSR and either 50ng pcDNA3.1 or 3xFLAG-MRAP2. Following 48-hours, media was changed to 50μl Hank’s buffered saline solution (HBSS), before loading each well with 40µL NanoGlo substrate (1:100 dilution, Promega) and luminescence read on a Glomax plate reader at 37°C. Vehicle and agonists were prepared in HBSS at 10x concentration and added to wells following recording of baseline signals for four cycles (equivalent to 8 minutes), then responses read for a further 22 minutes. The average baseline value recorded prior to agonist stimulation was subtracted from the experimental signal, then data normalized to vehicle-treated cells. AUC was calculated in GraphPad Prism and these values used to plot concentration-response curves with a 4-parameter sigmoidal fit.

### Glosensor cAMP assays

AdHEK cells were plated in 96-well plates and transfected with 100ng pGloSensor-20F, 50ng receptor (HA-SNAP-GHSR or HA-Halo-GHSR, HA-Halo-MC4R, HA-Halo-MC3R) and either 50ng pcDNA3.1 or 50ng 3xFLAG-MRAP2 (WT or mutant) plasmids per well. Cells were incubated for 48 hours then media changed to 40µL of equilibration media consisting of Fluorobrite DMEM (Invitrogen) containing a 2% (v/v) dilution of the GloSensor cAMP Reagent stock solution (Promega) with 1 µM IBMX (Sigma). Cells were incubated for 2 h at 37°C. For studies of GHSR, 10 µM forskolin (Sigma) was added to cells 10 minutes prior to recordings. Basal luminescence was read on a Glomax plate reader for 8 min, then MK0677 or NDP-MSH (5x concentration) with 1 µM IBMX added, and plates read for a further 30 minutes. Data was plotted in GraphPad Prism, area-within the curve calculated and values normalized to those at 0 µM agonist before plotting concentration-response curves with a 4-parameter sigmoidal fit.

### HILO

HILO imaging was performed as previously described^56,61^. AdHEK cells were seeded on 24mm coverslips (VWR) and transfected with 500ng of each plasmid 24-hours prior to experiments. For studies of HA-SNAP-GHSR, cells were cotransfected with 500 ng Venus-Kras as a plasma membrane marker. Cells were washed with imaging medium (Ca^2+^-and Mg^2+^-free HBSS with 10mM HEPES), then labelled with 2 μM of cell-membrane impermeable dyes (CLIP-surface 547 (BC-DY547, NEB) for FLAG-CLIP tagged plasmids or SNAP-surface-647 (NEB, Catalog# S91365) for HA-SNAP-GHSR tagged plasmids), or cell permeable dye (HaloTag R110Direct Ligand (Promega, Catalog#G3221) for HA-Halo tagged plasmids) in imaging medium for 20 minutes. Cells were washed and coverslips mounted onto plastic imaging chambers with a rubber seal and filled with imaging medium. HILO images were acquired on a custom-built TIRF microscope (Cairn Research) comprising an Eclipse Ti2 (Nikon) equipped with an EMCCD camera (iXon Ultra, Andor), a 488 nm diode laser, a hardware Perfect Focus System, a TIRF iLas2 module, and a 100× oil-immersion objective (NA 1.49, Nikon). The objective and samples were maintained at 37°C in a heated enclosure. Images were acquired on MetaMorph software (Molecular Devices) using a frame exposure of 50–200 ms with an image acquired before ligand stimulation (10 µM MK-0677) and a subsequent image taken every 30s thereafter, up to 20 minutes.

### IP-1 HTRF assays

IP-1 assays were performed with the Cisbio IP-One Gq HTRF kit (Revvity) as previously described^61,62^. Cells were co-transfected with 500 ng of GHSR constructs and 500 ng of either pcDNA3.1 or FLAG-MRAP2-WT. Forty-eight hours after transfection, cells were washed once in 1X PBS and resuspended in FluoroBrite DMEM (Thermo Fisher Scientific) supplemented with 10% FBS and 100 mM lithium chloride at 500 µL/well. Cells were then replated into 384-well plates at 7 µL/ well. MK-0677 agonist dilutions were made in stimulation buffer provided with the IP-One kit. MK-0677 was added to cells at 7 µL/well and incubated at 37°C for 1 hour. Cells were lysed according to the IP-One kit protocol and signal measured on a BMG Labtech PHERAstar microplate reader. The supplied IP-1 standard concentration range was run in tandem with each experimental repeat. Raw values were interpolated from the IP-1 standard curve and replotted as IP-1 concentration produced against agonist concentration. Non-linear log regression fit was performed using GraphPad Prism. Independent experiments were converted into a combined concentration-response graph by normalizing data to the maximal response of GHSR co-transfected with pcDNA3.1.

### Three-dimensional modelling of GHSR and MRAP2

Modeling of GHSR and MRAP2 was performed by AlphaFold2 using the ColabFold v1.5.2-patch in Google Co-laboratory^63^ and visualized using Pymol. FASTA sequences were obtained from Uniprot. Five models were predicted and the top ranked structure shown based on predicted local distance difference test (pLDDT).

### Assessment of cell surface expression

For assessment of GHSR surface expression, cells were transfected with 100ng HA-HALO-GHSR and 100 ng pcDNA or FLAG-MRAP2 (wild-type of mutant) and cells fixed 48-hours later in 4% PFA (Fisher Scientific UK) in PBS, then labelled with 1:1000 anti-HA mouse monoclonal antibody (BioLegend Cat#901514, RRID:AB_2565336) followed by Alexa Fluor 647 donkey anti-mouse secondary antibody (abcam Cat# ab181292, RRID:AB_3351687). Cells were washed, then fluorescence read on a Glomax plate reader. Data was normalized to that observed in cells transfected with pcDNA, set as 100.

## Statistical analysis

Statistical tests used for each experiment are indicated in the legends of each figure and the number of experimental replicates denoted by N. Data was plotted and statistical analyses performed in Graphpad Prism 7 or 9. Normality tests (Shapiro-Wilk or D’Agostino-Pearson) were performed on all datasets to determine whether parametric or non-parametric statistical tests were appropriate. A p value of <0.05 was considered statistically significant.

## Acknowledgements

Funding:

An Academy of Medical Sciences Springboard Award supported by the British Heart Foundation, Diabetes UK, the Global Challenges Research Fund, the Government Department of Business, Energy and Industrial Strategy and the Wellcome Trust. Ref: SBF004|1034 (C. Gorvin).

A Sir Henry Dale Fellowship jointly funded by the Wellcome Trust and the Royal Society. Grant Number 224155/Z/21/Z (C. Gorvin).

An NIH grant R01NS129904, the Rohr Family Research Scholar Award, and the Monique Weill-Caulier Award (J. Levitz).

The European Union’s Horizon Europe Framework Programme (deuterON, grant agreement number 101042046 (J. Broichhagen).

## Author contributions

Conceptualization: CMG

Methodology: CMG, JL, JB

Investigation: AJ, RAW, CMG

Writing – original draft: CMG

Writing – review and editing: All authors

## Competing interests

The authors declare no competing interests.

## Data and materials availability

All data needed to evaluate the conclusions in the paper are present in the paper and/or the Supplementary Materials. Plasmid constructs developed for this manuscript will readily be made available upon request. Plasmid constructs obtained from other researchers may be subject to Material Transfer Agreements. Please contact the corresponding author of this manuscript, or the named source for details.

## Notes

### Competing Interest Statement

The authors have declared no competing interest.

