## Supplementary Appendix for "MRAP2 potentiates GPCR signaling by conserved mechanisms that are disrupted by obesity-associated genetic variants"

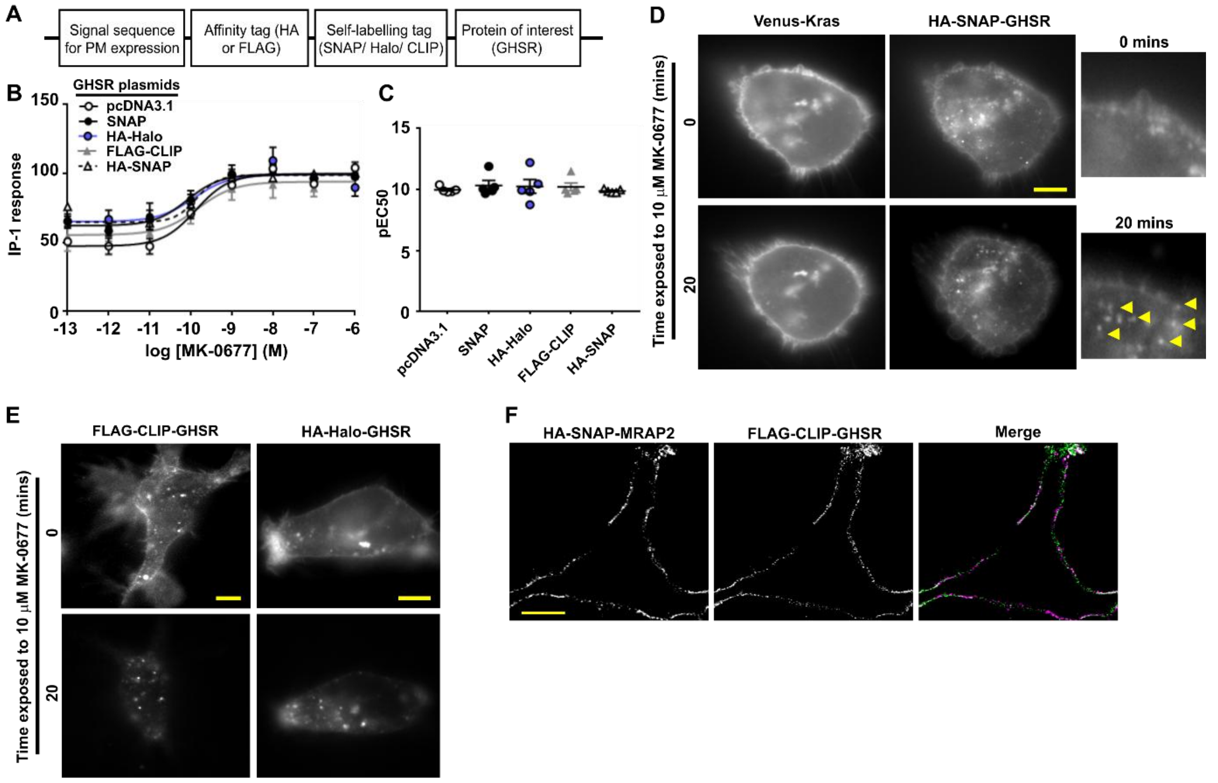


**Figure S1 GHSR plasmids signal and traffic normally**

(**A**) Schematic showing construction of the GHSR plasmids with a signal sequence, affinity tag (HA or FLAG), self-labeling tag (SNAP, Halo or CLIP) and the protein of interest. (**B**) GHSR-induced IP-1 activity in cells transfected with the three GHSR plasmids, pcDNA-3.1-GHSR and the commercially available SNAP-GHSR plasmid and (**C**) pEC50. N=5. The three plasmids had no differences in responses. (**D**) HILO images showing colocalization between the plasma membrane marker Venus-Kras and HA-SNAP-GHSR. Internalization of GHSR can be observed as an increased number of internalized vesicles (shown by arrows in close view) after 20-minute incubation with the MK-0677 agonist. (**E**) HILO images showing FLAG-CLIP-GHSR and HA-Halo-GHSR internalize normally. (**F**) SIM images showing colocalization of HA-SNAP-MRAP2 and FLAG-CLIP-GHSR at the cell surface. Scale, 5 μm.

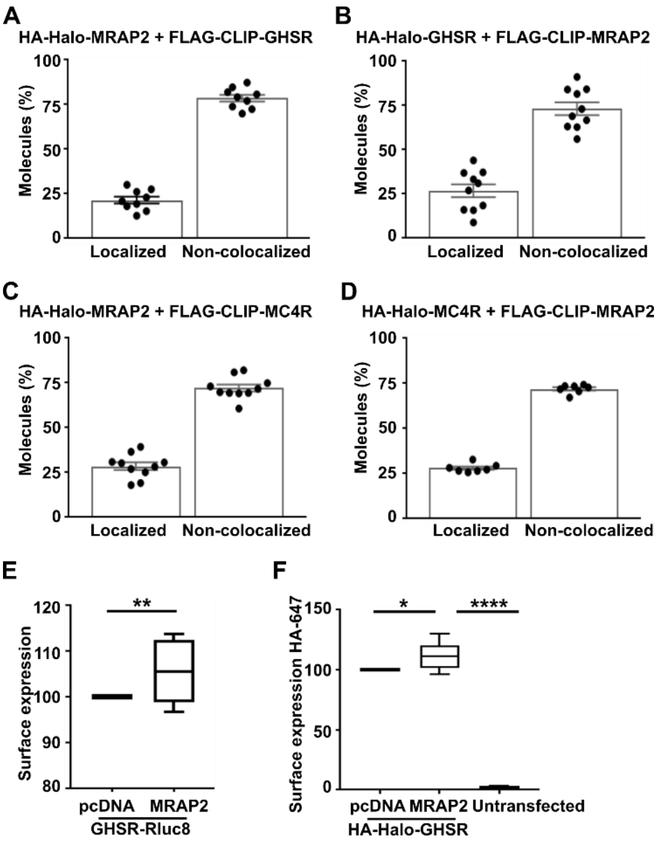


**Figure S2 Proportion of colocalized molecules in SiMPull analyses and surface expression of GHSR**

Proportion of molecules in two-color SiMPull that are colocalized in cells transfected with (**A**) HA-Halo-MRAP2 and FLAG-CLIP-GHSR, (**B**) HA-Halo-GHSR and FLAG-CLIP-MRAP2, (**C**) HA-Halo-MRAP2 and FLAG-CLIP-MC4R, and (**D**) HA-Halo-MC4R and FLAG-CLIP-MRAP2. (**E**) Assessment of GHSR-Rluc8 cell surface expression normalized to Kras as a cell surface marker in cells co-transfected with pcDNA or MRAP2. (**F**) Assessment of HA-Halo-GHSR cell surface expression measured by labelling surface GHSR with HA and a secondary Alexa Fluor 647 antibody. Fluorescence was normalized to untransfected cells. Statistical analyses were performed by unpaired t-test in E and one-way ANOVA with Dunnett's multiple-comparisons test in F.

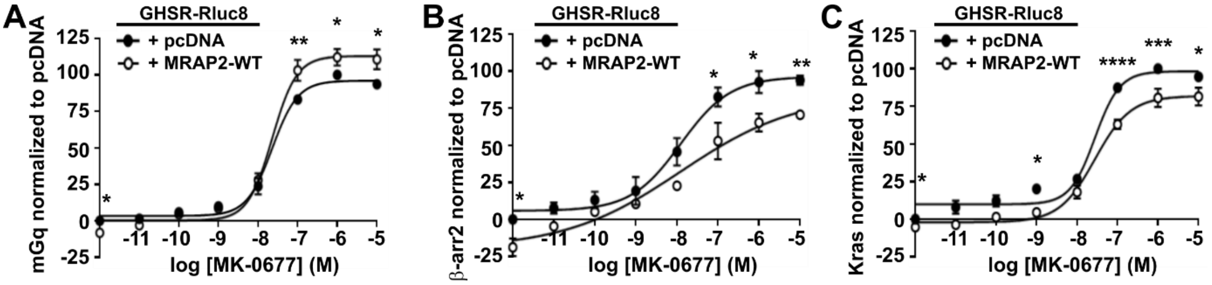


**Figure S3 MRAP2 enhances GHSR signaling and reduces internalization in mHypo-N39 mouse hypothalamic cell-lines**

(**A-C**) Normalized BRET responses measured between GHSR-Rluc8 and (A) mGq, (B) β-arrestin-2-YFP and (C) Venus-Kras in mHypo-N39 cells expressing MRAP2 or pcDNA. N = 3 biological replicates. Statistical analyses were performed by two-way ANOVA with Sidak's multiple-comparisons test. Statistics for constitutive activity, Emax and pEC50 are shown in Table S2. ****$p<0.0001$, ***$p<0.001$, **$p<0.01$, *$p<0.05$.

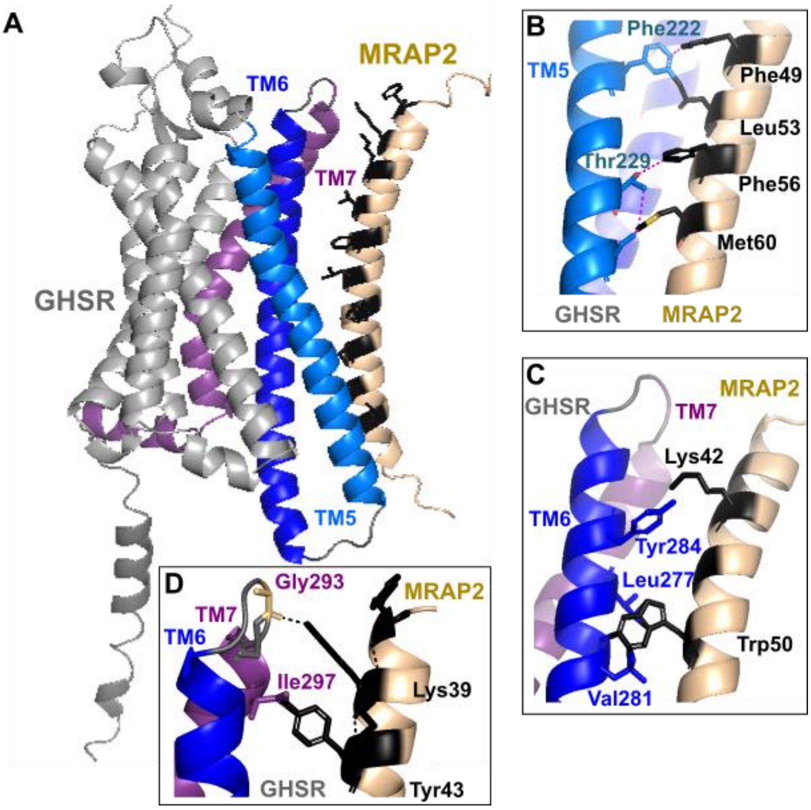


**Figure S4 Prediction of interaction between GHSR and MRAP2 transmembrane regions**

AlphaFold2 did not predict a consistent interaction interface with MRAP2 across all models. However, the model with the highest prediction score showed some interaction between MRAP2 and GHSR TM5, TM6 and the TM7-ECL3 region. (**A**) Highest ranked AlphaFold2 structural model of GHSR and MRAP2 in a 1-to-1 configuration with TM1-TM4 in gray and TM5, TM6 and TM7 highlighted in light blue, dark blue and purple respectively. (**B-D**) Predicted interactions between the MRAP2 transmembrane helix and residues of (B) TM5, (C) TM6 and (D) TM7 and ECL3 of GHSR. Predicted contacts are shown by dotted lines.

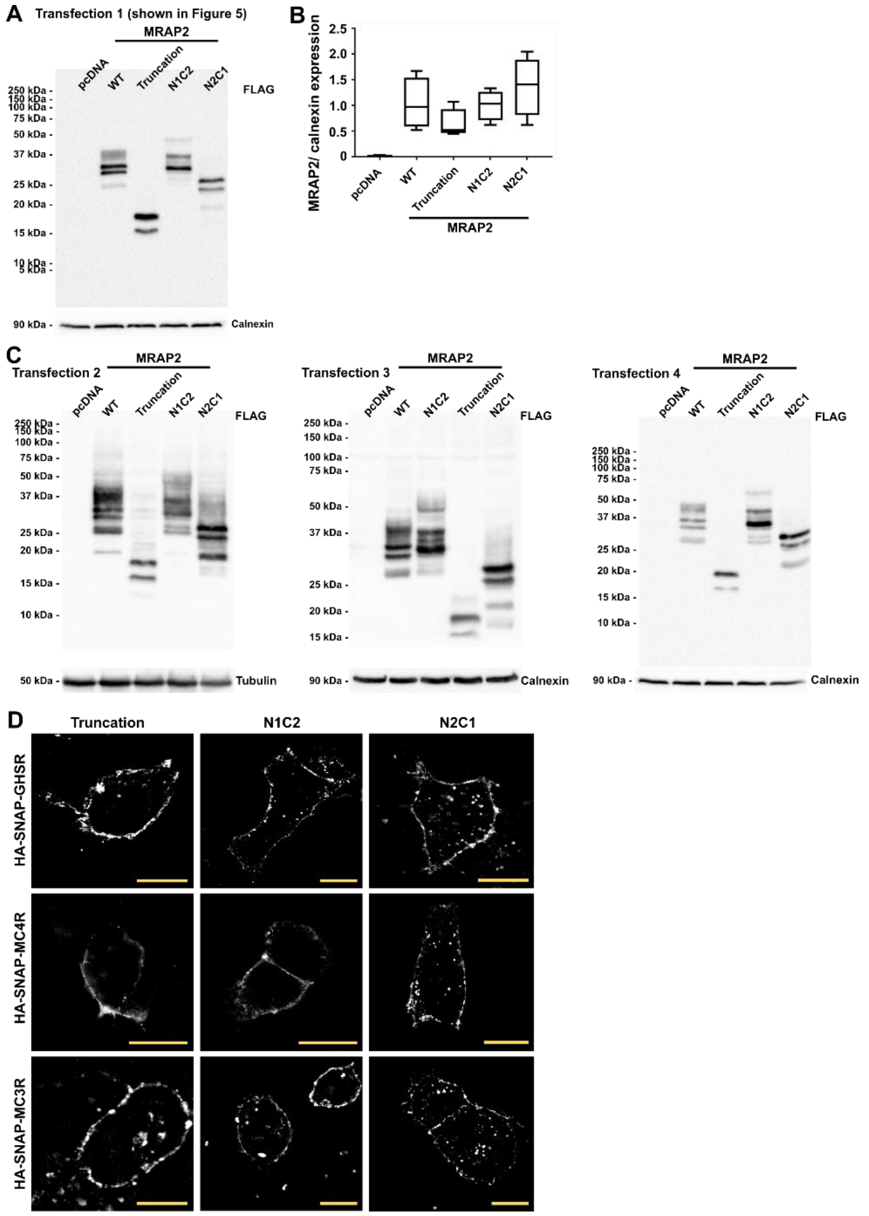


**Figure S5** **MRAP2 Truncation and chimeric mutants do not affect GPCR expression**

(**A**) Full blot from the image shown in Figure 5 of the FLAG-MRAP2 protein. Calnexin was used as a loading control. (**B**) Densitometry and (**C**) the full blots from four transfections of cells expressing either pcDNA, MRAP2-WT, -Truncation, -N1C2 or -N2C1. The amount of FLAG-MRAP2 was normalized to calnexin or tubulin loading controls and expressed as a fold-change relative to pcDNA expressing cells. Statistical analyses were performed by one-way ANOVA and showed no significant differences compared to MRAP2-WT. (**D**) SIM images showing HA-SNAP-GHSR, HA-SNAP-MC4R and HA-SNAP-MC3R are expressed at the surface of cells transfected with MRAP2-Truncation, MRAP2-N1C2 and MRAP2-N2C1 plasmids. Scale, 5 μm.

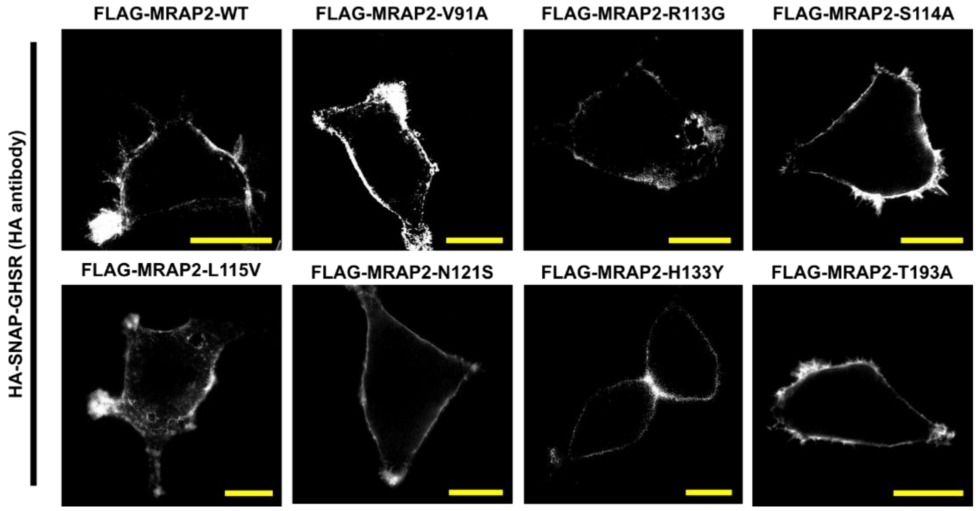


**Figure S6 GHSR expression in the presence of MRAP2 human variants**
GHSR surface expression in non-permeabilized cells transfected with the MRAP2 human variants, measured by SIM. Scale, 5 µm.

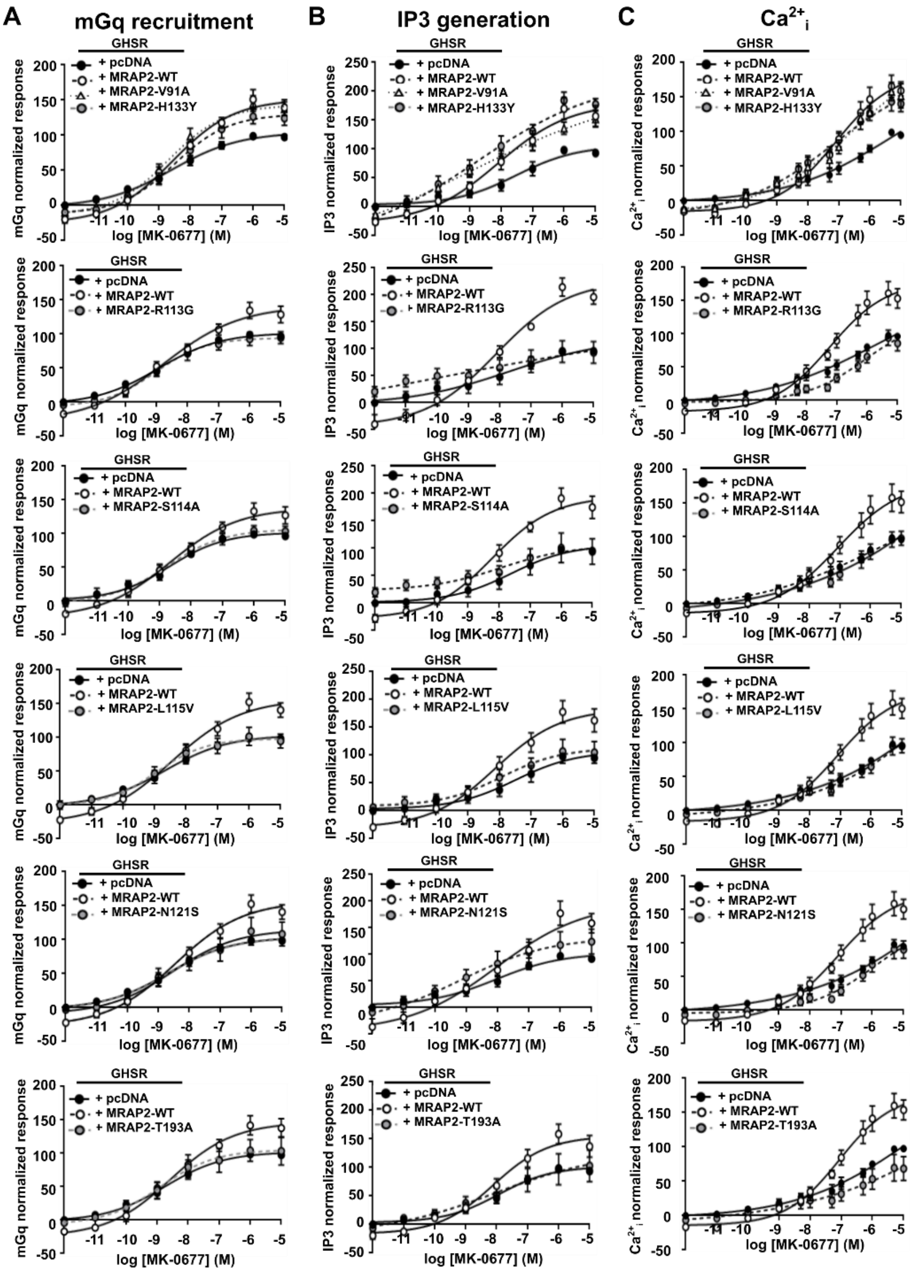


**Figure S7 Effect of MRAP2 obesity-associated variants on GHSR-mediated Gq/11 signaling** (**A**) Normalized BRET responses measured between GHSR-Rluc8 and Venus-mGq in cells expressing either pcDNA, MRAP2-wild type (WT) or MRAP2 variants. Basal activity and maximal responses with statistical analyses are shown in Figure 7A-B. N = 8 for R113G, S114A, N=6 for V91A, T193A, N = 7 for L115V, N121S, H133Y. (**B**) Normalized IP3 responses measured by IP3 NanoBiT biosensor in cells expressing HA-Halo-GHSR with either pcDNA, MRAP2-wild type (WT) or MRAP2 mutants. N = 7 for N88Y, V91A, N121S, N = 6 for R113G, S114A, L115V, H133Y, T193A. Basal activity and maximal responses with statistical analyses are shown in Figure 7C-D. AUC was measured and responses expressed relative to the pcDNA maximal response set at 100 in A and B. (**C**) Normalized $Ca^{2+}_{i}$ responses measured by Fluo-4 assays in cells expressing HA-Halo-GHSR with either pcDNA, MRAP2-wild type (WT) or MRAP2 mutants. Maximal responses at each concentration were measured and responses expressed relative to the pcDNA maximal response set at 100. N = 8 for N88Y, R113G, L115V, N121S, H133Y, T193A, N = 7 for V91A and S114A. Basal activity and maximal responses with statistical analyses are shown in Figure 7E-F. Data shows mean±SEM. V91A and H133Y have been reported in normal weight individuals, other variants have been associated with overweight and/or obesity. pEC50 values are shown in Table S6.

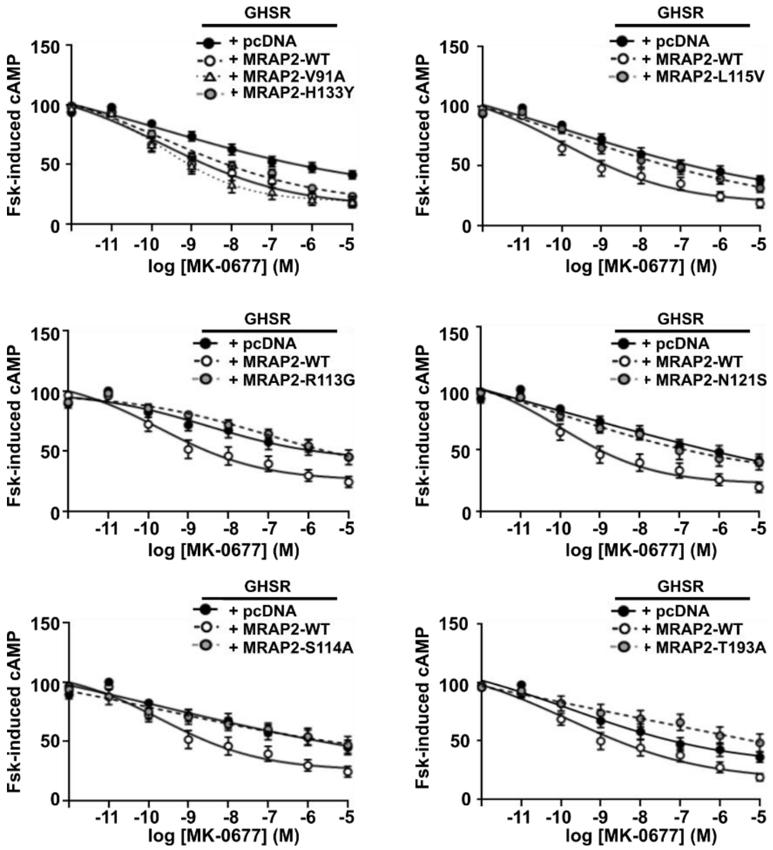


**Figure S8 Effect of MRAP2 obesity-associated variants on GHSR-mediated Gi/o signaling**

Suppression of forskolin-induced increases in cAMP, measured by cAMP Glosensor in cells expressing HA-Halo-GHSR with either pcDNA, MRAP2-wild type (WT) or MRAP2 variants. AUC was measured and responses expressed relative to the pcDNA maximal response set at 100. Maximal responses are shown in Figure 7G. Data shows mean±SEM. N = 10 for V91A, L115V, H133Y, N = 9 for T193A, N = 8 for N121S, N = 7 for R113G, S114A. V91A and H133Y have been reported in normal weight individuals, other variants have been associated with overweight and/or obesity. pEC50 values are shown in Table S6.

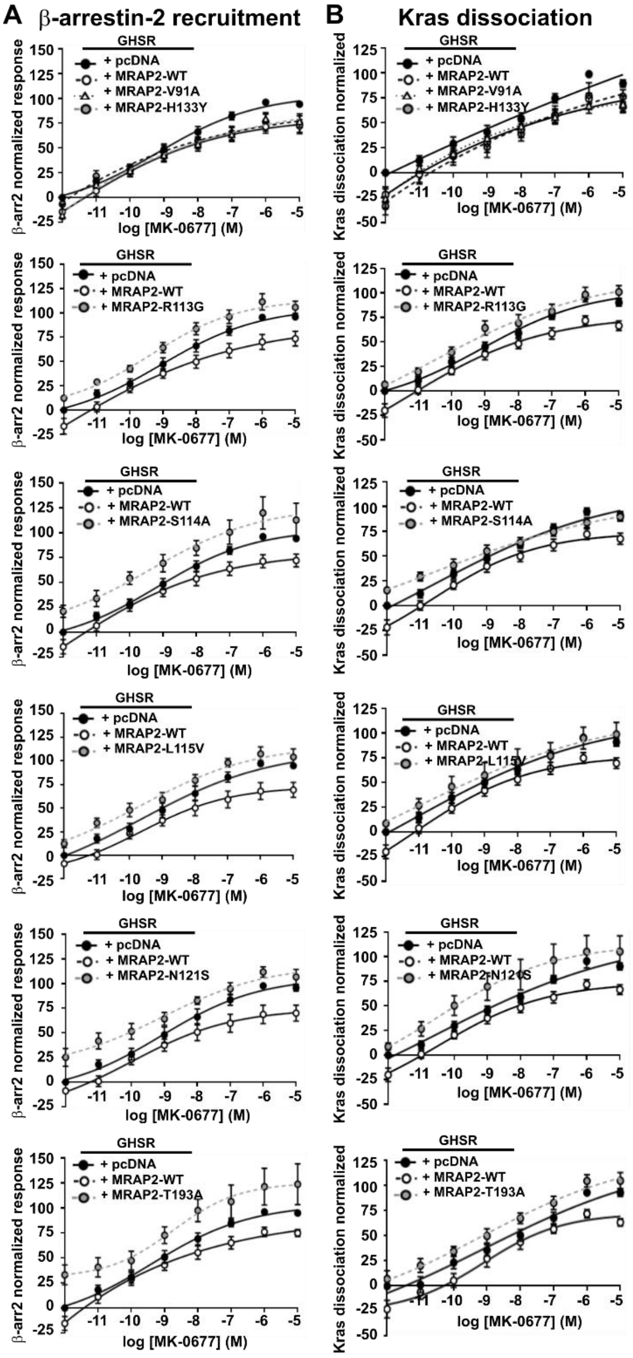


**Figure S9 Effect of MRAP2 obesity-associated variants on GHSR-mediated trafficking**

(**A-B**) Normalized BRET responses measured between GHSR-Rluc8 and (A) β-arr2-YFP and (B) Venus-Kras in cells expressing either pcDNA, MRAP2-wild type (WT) or MRAP2 mutants. AUC was measured and responses expressed relative to the pcDNA maximal response set at 100. Data shows mean±SEM. N = 7 for S114A, N121S, H133Y, T193A, N = 6 for V91A, R113G, N = 5 for L115V for A. N = 8 for V91A, R113G, L115V, N121S, H133Y, T193A, N = 7 for S114A for B. V91A and H133Y have been reported in normal weight individuals, other variants have been associated with overweight and/or obesity. pEC50 values are shown in Table S6 and maximal responses and constitutive activity comparisons shown in Figure 7.

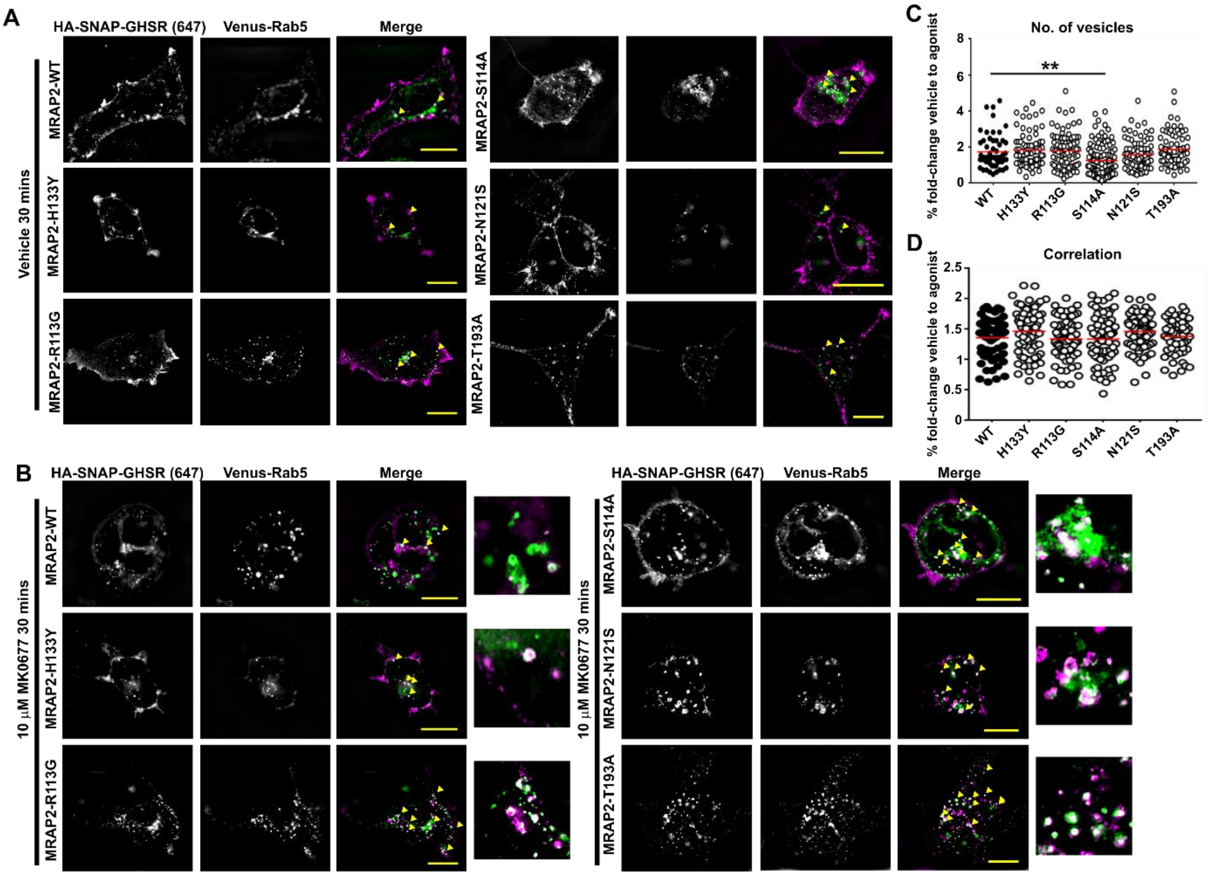


**Figure S10 Effect of MRAP2 obesity-associated variants on GHSR-mediated constitutive internalization**

(**A**) Control SIM images of HA-SNAP-GHSR and Venus-Rab5 in cells expressing either pcDNA, MRAP2-wild type (WT) or MRAP2 mutants and exposed to vehicle for 30 minutes. Quantification of the number of internalized vesicles and colocalization is shown in Figure 8. The number of cells quantified from six independent transfections were as follows: WT vehicle (64), H133Y vehicle (65), R113G vehicle (76), S114A vehicle (88), N121S vehicle (77), T193A vehicle (65). Yellow arrows show areas of colocalization. (**B**) SIM images of HA-SNAP-GHSR and Venus-Rab5 in cells expressing either pcDNA, MRAP2-wild type (WT) or MRAP2 mutants following exposure of cells to the GHSR agonist MK-0677 for 30 minutes, with zoomed image shown to the right. The number of cells quantified from six independent transfections were as follows: WT vehicle (64) and agonist (64), H133Y vehicle (65) and agonist (74), R113G vehicle (76) and agonist (71), S114A vehicle (88) and agonist (78), N121S vehicle (77) and agonist (81), T193A vehicle (65) and agonist (58). Yellow arrows show areas of colocalization. Scale, 5 μm. MRAP2-H133Y has only been identified in normal weight individuals, while other variants have been reported in overweight or obesity. (**C-D**) Percentage fold-change difference in internalization following treatment with vehicle or MK0677. Fold-change responses were compared to the mean internalization in each group normalized to 1. Fold-change differences were calculated from (C) number of vesicles data in Figure 7O and (D) colocalization data in Figure 7P. Statistical analyses were performed by one-way ANOVA with Dunnett's multiple-comparisons test. **$p<0.001$.

**Table S1 Expression plasmids used in this manuscript**

| Plasmid name | Information | Source |
|---|---|---|
| ss-HA-Halo-GHSR | N-terminal signal peptide from mGluR5, followed by HA, HALO and human GHSR | This manuscript |
| ss-HA-SNAP-GHSR | N-terminal signal peptide from mGluR5, followed by HA, SNAP and human GHSR | Lead contact, Caroline Gorvin, University of Birmingham [43] |
| ss-FLAG-CLIP-GHSR | N-terminal signal peptide from mGluR5, followed by FLAG, CLIP and human GHSR | This manuscript |
| ss-HA-Halo-MRAP2 | N-terminal signal peptide from mGluR5, followed by HA, Halo and human MRAP2 | Lead contact, Caroline Gorvin, University of Birmingham [8] |
| ss-HA-SNAP-MRAP2 | N-terminal signal peptide from mGluR5, followed by HA, SNAP and human MRAP2 | Lead contact, Caroline Gorvin, University of Birmingham [8] |
| ss-FLAG-CLIP-MRAP2 | N-terminal signal peptide from mGluR5, followed by FLAG, CLIP and human MRAP2 | Lead contact, Caroline Gorvin, University of Birmingham [8] |
| ss-HA-Halo-MC4R | N-terminal signal peptide from mGluR5, followed by HA, Halo and human MC4R | Lead contact, Caroline Gorvin, University of Birmingham [17] |
| ss-FLAG-CLIP-MC4R | N-terminal signal peptide from mGluR5, followed by FLAG, CLIP and human MC4R | Lead contact, Caroline Gorvin, University of Birmingham [17] |
| ss-HA-SNAP-MC4R | N-terminal signal peptide from mGluR5, followed by HA, SNAP and human MC4R | Lead contact, Caroline Gorvin, University of Birmingham [17] |
| ss-HA-Halo-MC3R | N-terminal signal peptide from mGluR5, followed by HA, Halo and human MC3R | Lead contact, Caroline Gorvin, University of Birmingham [8] |
| ss-FLAG-CLIP-MC3R | N-terminal signal peptide from mGluR5, followed by FLAG, CLIP and human MC3R | Lead contact, Caroline Gorvin, University of Birmingham [8] |
| ss-HA-SNAP-MC3R | N-terminal signal peptide from mGluR5, followed by HA, SNAP and human MC3R | Lead contact, Caroline Gorvin, University of Birmingham [8] |
| ss-HA-Halo-mGluR2 | Used as template for ss-HA-Halo-GHSR | Joshua Levitz, Weill Cornell Medicine |
| ss-FLAG-CLIP-mGluR2 | Used as template for ss-FLAG-CLIP-GHSR | Joshua Levitz, Weill Cornell Medicine |
| SNAP-GHSR | Used as template for GHSR Halo/ SNAP/ CLIP constructs and in test of constructs. | Cisbio |
| GHSR-LgC | NanoBiT interaction assays. C-terminal LgC tag. | This manuscript |
| MRAP2-SmC | NanoBiT interaction assays. C-terminal SmC tag. | Lead contact, Caroline Gorvin, University of Birmingham [8] |
| pBiT2.1 C [TK/SmBiT] | NanoBiT interaction assays | Promega |
| pBiT1.1 C [TK/LgBiT] | NanoBiT interaction assays | Promega |
| cAMP Glosensor-22F | cAMP sensor | Promega |
| LgBiT-IP3R2-SmBiT | IP$_3$ biosensor | Asuka Inoue, Tohoku University [48] |
| MRAP2-3xFLAG | MRAP2 with C-terminal 3x FLAG | Julien Sebag, University of Iowa |
| B-arrestin-2-mYFP | BRET | Robert Lefkowitz, Addgene plasmid #36917 |
| pcDNA3.1-GHSR | Used as control in IP-1 assays | This manuscript |
| GHSR-Rluc8 | BRET | Lead contact, Caroline Gorvin, University of Birmingham [39] |
| Venus-mGq | BRET | Nevin Lambert, Augusta University [67] |
| Venus-Kras | BRET/SIM | Nevin Lambert, Augusta University [67] |
| Venus-Rab5 | SIM | Nevin Lambert, Augusta University [67] |
| MRAP2-Truncation | N-terminal HA tag, MRAP2 1-63, 3xFLAG tag | This manuscript |
| MRAP2-N1C2 | N-terminal HA tag, MRAP1 1-56, MRAP2 64-205, 3xFLAG tag | This manuscript |
| MRAP2-N2C1 | N-terminal HA tag, MRAP2 1-63, MRAP1 57-172, 3xFLAG tag | This manuscript |
| MRAP2-3xFLAG-V91A | BRET, Glosensor, fluo-4, SIM | Lead contact, Caroline Gorvin, University of Birmingham [17] |
| MRAP2-3xFLAG-R113G | BRET, Glosensor, fluo-4, SIM | Lead contact, Caroline Gorvin, University of Birmingham [17] |
| MRAP2-3xFLAG-S114A | BRET, Glosensor, fluo-4, SIM | Lead contact, Caroline Gorvin, University of Birmingham [17] |
| MRAP2-3xFLAG-L115V | BRET, Glosensor, fluo-4, SIM | Lead contact, Caroline Gorvin, University of Birmingham [17] |
| MRAP2-3xFLAG-N121S | BRET, Glosensor, fluo-4, SIM | Lead contact, Caroline Gorvin, University of Birmingham [17] |
| MRAP2-3xFLAG-H133Y | BRET, Glosensor, fluo-4, SIM | Lead contact, Caroline Gorvin, University of Birmingham [17] |
| MRAP2-3xFLAG-T193A | BRET, Glosensor, fluo-4, SIM | Lead contact, Caroline Gorvin, University of Birmingham [17] |

**Table S2 Effect of MRAP2 on GHSR constitutive activity, maximal responses and pEC50 values**

| Cell-line | Assay | Comparison | pcDNA | MRAP2 | p-value |
|---|---|---|---|---|---|
| **AdHEK** | **IP3 luminescence** (N = 11) | Constitutive activity | 0 | -28.31 ± 10.62 | * |
| | | Emax | 100 | 178.7 ± 15.84 | **** |
| | | pEC50 | 8.37 ± 0.43 | 8.06 ± 0.19 | ns |
| | **$Ca^{2+}_i$ (Fluo-4)** (N=9) | Constitutive activity | 0 | -16.6 ± 2.01 | **** |
| | | Emax | 100 | 162.5 ± 15.74 | ** |
| | | pEC50 | 6.22 ± 0.35 | 6.42 ± 0.20 | ns |
| | **mGq BRET** (N=10) | Constitutive activity | 0 | -20.6 ± 3.62 | **** |
| | | Emax | 100 | 139.4 ± 11.24 | ** |
| | | pEC50 | 8.55 ± 0.23 | 8.67 ± 0.27 | ns |
| | **cAMP suppression** (N=12) | Constitutive activity | N/A | N/A | |
| | | Emax | 39.52 ± 3.76 | 19.64 ± 2.96 | ** |
| | | pEC50 | 8.45 ± 0.34 | 9.24 ± 0.40 | ns |
| | **β-arrestin-2 recruitment** (N=10) | Constitutive activity | 0 | -11.86 ± 5.14 | * |
| | | Emax | 100 | 76.47 ± 4.56 | **** |
| | | pEC50 | 8.89 ± 0.19 | 10.09 ± 0.30 | ** |
| | **Kras dissociation** (N=13) | Constitutive activity | 0 | -24.04 ± 5.58 | *** |
| | | Emax | 100 | 77.23 ± 3.69 | **** |
| | | pEC50 | 8.53 ± 0.26 | 9.50 ± 0.24 | * |
| **mHypo-N39** | **mGq BRET** (N=3) | Constitutive activity | 0 | -8.91 ± 2.56 | * |
| | | Emax | 100 | 114.6 ± 4.72 | * |
| | | pEC50 | 8.58 ± 0.09 | 8.70 ± 0.04 | ns |
| | **β-arrestin-2 recruitment** (N=3) | Constitutive activity | 0 | -20.51 ± 7.64 | * |
| | | Emax | 100 | 71.31 ± 2.35 | *** |
| | | pEC50 | 8.84 ± 0.24 | 8.97 ± 0.09 | ns |
| | **Kras dissociation** (N=3) | Constitutive activity | 0 | -6.03 ± 0.71 | ** |
| | | Emax | 100 | 81.41 ± 5.96 | * |
| | | pEC50 | 8.59 ± 0.04 | 8.53 ± 0.18 | ns |

Statistical analyses from concentration-response curves shown in Figure 3 and S3. Constitutive activity cannot be calculated for cAMP as responses are relative to induction of cAMP by forskolin, which may be variable between samples. Therefore, these responses are normalized to the forskolin-induced cAMP response at 0 M MK-0677 for each condition. Statistical analyses were performed by unpaired t-test. ns, not significant. ****p<0.0001, ***p<0.001, **p<0.01, *p<0.05.

**Table S3 Effect of LEAP2 on pEC50 values**

| LEAP2 (nM) | pcDNA | MRAP2-WT |
|---|---|---|
| 0 | 8.44 ± 0.16 | 8.04 ± 0.10 |
| 1 | 8.02 ± 0.52 | 7.40 ± 0.23 |
| 10 | 8.46 ± 0.55 | 6.93 ± 0.35 |
| 100 | 8.63 ± 0.87 | 7.61 ± 0.51 |

pEC50 from concentration-response curves shown in Figure 3. Statistical analyses were performed by one-way ANOVA with Sidak's multiple-comparisons test and showed no significant differences.

**Table S4 Effect of GHSR transmembrane interfering peptides on pEC50 values**

| Interfering peptide | pcDNA | MRAP2-WT |
|---|---|---|
| None | 8.39 ± 0.30 | 8.26 ± 0.35 |
| All | 7.39 ± 0.31 | 7.29 ± 0.47 |
| TM1 | 8.38 ± 0.34 | 8.44 ± 0.57 |
| TM2 | 7.99± 0.25 | 8.23 ± 0.36 |
| TM3 | 7.94 ± 0.26 | 8.13 ± 0.31 |
| TM4 | 8.65 ± 0.22 | 8.58 ± 0.10 |
| TM5 | 8.47 ± 0.29 | 8.25 ± 0.28 |
| TM6 | 7.82 ± 0.45 | 7.32 ± 0.37 |
| TM7 | 7.77 ± 0.35 | 7.47 ± 0.20 |

pEC50 from concentration-response curves shown in Figure 4. Statistical analyses were performed by one-way ANOVA with Sidak's multiple-comparisons test and showed no significant differences.

**Table S5 Effect of MRAP2 chimaeras and truncation mutants on pEC50 values**

| | | GHSR | | | | | MC4R | MC3R |
|---|---|---|---|---|---|---|---|---|
| | **mGq BRET** | **$Ca^{2+}_i$** | **Suppression of cAMP** | **β-arrestin-2 recruitment** | **Kras dissociation** | | **cAMP** | **cAMP** |
| **pcDNA** | 7.44 ± 0.22 | 6.71 ± 0.34 | 8.53 ± 0.54 | 7.55 ± 0.17 | 7.85 ± 0.25 | | 7.28 ± 0.17 | 7.31 ± 0.12 |
| **MRAP2-WT** | 7.52 ± 0.56 | 6.98 ± 0.25 | 9.21 ± 0.30 | 7.75 ± 0.30 | 8.32 ± 0.32 | | 7.49 ± 0.21 | 7.24 ± 0.10 |
| **MRAP-2-Truncation** | 7.56 ± 0.28 | 7.05 ± 0.15 | 8.16 ± 0.59 | 7.84 ± 0.31 | 8.12 ± 0.20 | | 7.77 ± 0.36 | 7.11 ± 0.11 |
| **MRAP2-N2C1** | 7.47 ± 0.87 | 7.04 ± 0.17 | 8.39 ± 0.67 | 7.41 ± 0.17 | 8.99 ± 0.25 | | 8.48 ± 0.62 | 7.34 ± 0.15 |
| **MRAP2-N1C2** | 7.46 ± 0.26 | 6.63 ± 0.34 | 8.36 ± 0.44 | 7.91 ± 0.35 | 7.85 ± 0.37 | | 7.28 ± 0.28 | 7.37 ± 0.14 |

pEC50 from concentration-response curves shown in Figure 5. Statistical analyses were performed by one-way ANOVA with Sidak's multiple-comparisons test and showed no significant differences when compared to pcDNA or MRAP2-WT.

**Table S6 Effect of MRAP2 human obesity-associated variants on GHSR pEC50 values**

| | **mGq recruitment** | **IP3** | **$Ca^{2+}_i$** | **Suppression of cAMP** | **β-arrestin-2 recruitment** | **Kras dissociation** |
|---|---|---|---|---|---|---|
| **pcDNA** | 8.37 ± 0.29 | 7.84 ± 0.48 | 6.22 ± 0.35 | 8.45 ± 0.34 | 8.83 ± 0.24 ** | 8.25 ± 0.29 * |
| **MRAP2-WT** | 8.76 ± 0.24 | 7.94 ± 0.30 | 6.42 ± 0.20 | 9.24 ± 0.40 | 10.01 ± 0.23 | 9.65 ± 0.36 |
| **V91A** | 8.68 ± 0.21 | 8.57 ± 0.49 | 6.40 ± 0.34 | 9.28 ± 0.37 | 9.81 ± 0.51 | 9.97 ± 0.43 |
| **H133Y** | 8.54 ± 0.30 | 8.63 ± 0.47 | 6.67 ± 0.29 | 9.50 ± 0.22 | 9.76 ± 0.29 | 9.42 ± 0.55 |
| **R113G** | 8.98 ± 0.23 | 8.74 ± 0.79 | 5.58 ± 0.15 | 8.20 ± 0.39 | 8.88 ± 0.10 * | 8.82 ± 0.47 |
| **S114A** | 8.87 ± 0.24 | 8.78 ± 0.71 | 5.83 ± 0.24 | 9.44 ± 0.59 | 8.81 ± 0.36 ** | 9.37 ± 0.45 |
| **L115V** | 8.89 ± 0.25 | 8.08 ± 0.17 | 5.94 ± 0.22 | 8.78 ± 0.46 | 8.95 ± 0.29 * | 8.34 ± 0.54 |
| **N121S** | 8.35 ± 0.40 | 8.54 ± 0.39 | 5.45 ± 0.07 | 8.92 ± 0.46 | 8.67 ± 0.14 ** | 9.22 ± 0.19 |
| **T193A** | 8.68 ± 0.23 | 8.51 ± 0.38 | 6.40 ± 0.39 | 8.88 ± 0.50 | 9.09 ± 0.25 * | 8.99 ± 0.34 |

pEC50 from concentration-response curves shown in Figure 7 and S6-S8. Statistical analyses were performed by one-way ANOVA with Sidak's multiple-comparisons test comparing responses to MRAP2-WT. V91A and H133Y have been reported in normal weight individuals, other variants have been associated with overweight and/or obesity.
